# Declines in mRNA synthesis set the rate of organismal aging

**DOI:** 10.64898/2026.07.02.733740

**Authors:** Andrea Del Carmen-Fabregat, Natasha Oswal, Kshitij Sinha, Jeremy Vicencio, Ryan Abramowitz, Matthias Eder, Oguzhan Begik, Lucia Sedlackova, Eva Maria Novoa, Nicholas Stroustrup

## Abstract

The abundance of mRNA sets a ceiling on a cell’s capacity to produce protein and carry out its functions. Here, we describe a pathological decline in absolute mRNA abundance that occurs in most cell types during invertebrate and mammalian aging, caused by decreases in mRNA synthesis capacity. In *C. elegans*, decreases in mRNA abundance are tightly coupled to declines in RNA Polymerase II (Pol II) protein abundance. Measuring Pol II abundance dynamics *in vivo*, we find that individuals enter adulthood with a four-fold excess of Pol II, whose kinetic equilibration towards its homeostatic set point drives reductions in mRNA abundance and, in turn, organismal aging. Briefly accelerating Pol II declines produces permanent, dose-dependent reductions in healthspan and lifespan, whereas deceleration extends both. Although disrupted cellular homeostasis is conventionally seen as a consequence of aging, our results reveal how an out-of-equilibrium state established during development provides a driving force for aging.

## INTRODUCTION

The mechanisms of gene-regulation provide cells, tissues, and organisms with a flexible means for maintaining physiologic homeostasis—coordinating the abundance of proteins and their downstream activities. However, during aging, the capacity to maintain homeostasis is lost in part due to widespread changes in gene regulation that include global length-dependent transcriptomic biases^1^ and altered mRNA processing^2^ and degradation^3^, which result in diverse pathological consequences^4^. Yet these pathological changes in global gene expression become apparent only in aged individuals, as consequences of aging—leaving unexplained the ultimate causes of gene-regulatory dysfunction that act early in adulthood to drive individuals out of youth and towards old age.

In this study, we combine absolute mRNA quantification methods and quantitative modelling of *in vivo* protein dynamics to identify a non-equilibrium cellular state in mRNA synthesis at the very beginning of adulthood. Previous reports have identified decreases in the number of RNA-seq reads mapping to certain cell types in aged samples^5–7^. Here, we find that decreases in mRNA abundance during aging are widespread across cell types in both *C. elegans* and mice. In *C. elegans*, we show such decreases are driven by declines in a cell’s capacity for mRNA synthesis, reflected in an age-dependent drop in RNA Polymerase II (Pol II) abundance in the nucleus during aging starting at the onset of adulthood. Experimental modulation of the rate of Pol II declines produces tight dose-dependent changes in the organismal aging rate, highlighting the causal role of mRNA synthesis declines in both cellular and organismal aging.

## RESULTS

### Global mRNA abundance of organisms and cells decreases during aging at a rate strongly correlated with lifespan

To study the dynamics of global changes in gene regulation during aging, we adapted a single-individual RNA-seq protocol^8^ for absolute quantification of organismal mRNA abundance (*See Methods, Suppl. Note 1*), defined as the total abundance of all mRNA transcripts in aggregate—a quantity sometimes referred to as “transcriptome size”^9^. We find that individual wild-type *C. elegans* nematodes contain on average 700 million molecules of mRNA in youth (day 2 of adulthood), and that this abundance drops two-fold by old age (day 9) and eight-fold by very old age (day 21) (Fig. 1a, Fig. S1a-c). This large-magnitude effect will be missed by any aging study that employs conventional RNA-seq library size normalization methods, which implicitly assume all samples have equal mRNA concentrations and rescale total transcript counts, masking global shifts in mRNA abundance.

**Figure 1.**
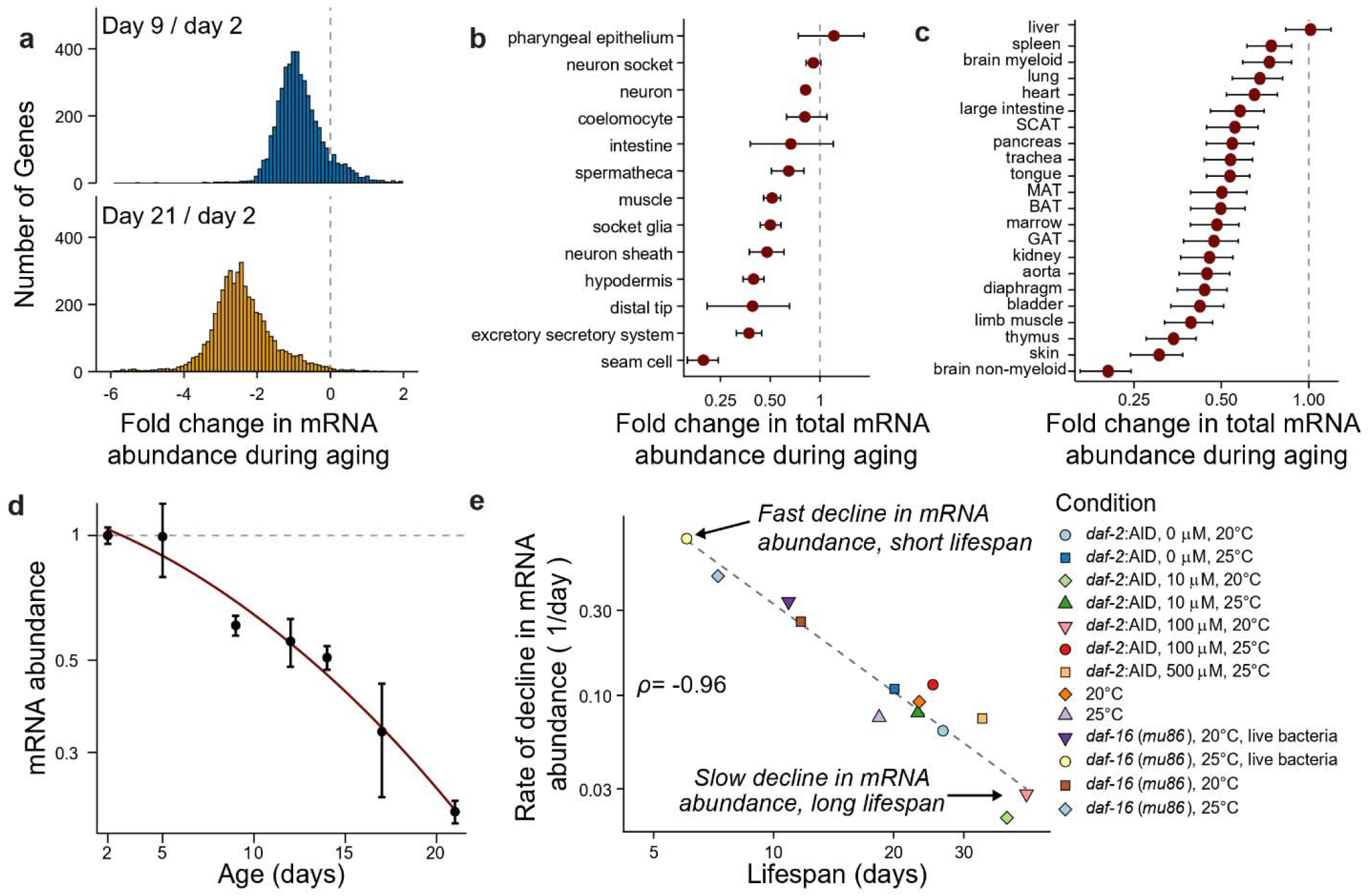
mRNA abundance of cells and organisms decreases during aging and the rate of decline closely tracks lifespan. **a**. Fold-change in the absolute abundance of each mRNA per individual among old (day 9) and very old (day 21) wild-type (N2) *C. elegans* relative to young (day 2). **b**. The mean change in total mRNA abundance per cell in day 11 compared to day 1 *C. elegans* grouped by cell-type in Roux *et al*. (2023). **c**. The mean change in total mRNA abundance per cell in 18 months compared to 3 months C57BL/6JN adult mice, grouped by tissue-of-origin in Tabula Muris Senis (2020). **d**. The progressive decline of total mRNA abundance across the lifespan of a wild-type *C. elegans* cohort relative to young (day 2) individuals. **e**. Relationship between the rate of decline in mRNA abundance during adulthood *(Statistical methods)* and lifespan across 13 lifespan-altering interventions in *C. elegans*, with linear regression line *(dashed)*, Pearson correlation = -0.96, P value <0.001. Panels *a, d* and *e*, show the mean of 4 biological replicates of n=30 each.

Decreases in organismal mRNA abundance are driven by a transcriptome-wide decrease in abundance and not by a decrease in a specific class of transcripts—leading to a scaling of the entire distribution of gene-expression (Fig. S1e). We similarly observe large declines in mRNA in the somatic tissues of germline-ablated *glp-1(e2141)* populations—distinct from any age-associated changes in germline structure or activity (Fig. S1d,e). The decrease in organismal mRNA is not attributable to an age-associated decrease in body size, as mRNA abundance decreases over a period in which body size increases and then decreases again (Fig. S1f,g). To confirm our findings using an orthogonal approach, we applied native mRNA long-read sequencing to characterize *glp-1(e2141)* populations and again identified a large decline in mRNA abundance occurring with age (Fig. S1h).

Multicellular organisms contain a variety of cell types that differ in mRNA abundance and transcriptome composition. Analyzing published single-cell and single-nucleus *C. elegans* transcriptomes^10,11^, we find that the age-associated declines in mRNA abundance occur across most cell-types (Fig. 1b, Fig. S1i,j). We conclude that in *C. elegans*, an age-dependent decrease in organismal mRNA levels results from the cumulative contribution of parallel decreases occurring within individual nuclei, cells, and tissues.

Next, we investigated whether such declines in mRNA abundance are evolutionarily conserved. Analysis of published single-cell murine transcriptomes^12^ revealed pervasive reductions in cellular mRNA abundance across most tissues and cell-types, with declines of more than two-fold in the aorta and more than three-fold in brain non-myeloid tissues by old age (18 and 24 months) relative to young adult mice (3 months) (Fig. 1c, Fig. S1k-m). These observations indicate that the progressive, disproportionate depletion of cellular and tissue mRNA levels is a conserved feature of aging across multi-cellular animals.

Declines in organismal and cellular mRNA abundance in *C. elegans* occur progressively throughout adulthood (Fig. 1d). To investigate whether these changes are coupled to aging—and not merely related to the passage of chronological time—we considered multiple lifespan-altering interventions including changes in food, temperature, *daf-16*/FOXO inactivation and grades of *daf-2* insulin/IGF-1 signaling modulation. We find that the rate of decline in global mRNA abundance is tightly coupled to a population’s lifespan (Fig. 1e, Fig. S1n), predicting lifespan with a Pearson correlation of 0.96. These results suggest an important link between declines in mRNA abundance and the organismal aging processes that determine lifespan— identifying global changes in mRNA abundance as a crucial hallmark of cellular and organismal aging.

### RNA Polymerase II decreases in abundance during aging, limiting global mRNA abundance and lifespan

Previous studies have shown that the nuclear abundance of Pol II can act as the main limiting factor for mRNA synthesis^13^. Therefore, we next investigated the physiologic role of Pol II abundance in determining transcriptional output. Because the Pol II subunit RPB-2 must be assembled into a full Pol II complex prior to nuclear localization, we developed RPB-2::GFP as a quantitative reporter for the nuclear abundance of the Pol II complex in live animals (Fig. 2a, *left*). We identify a global drop in the abundance of RPB-2 in aging (Fig. 2a,b) occurring in all somatic tissues but not in reproductive germline tissues (Fig. S2a), with RPB-2 levels declining at a slower rate in heads compared to the rest of the body (Fig. S2b).

**Figure 2.**
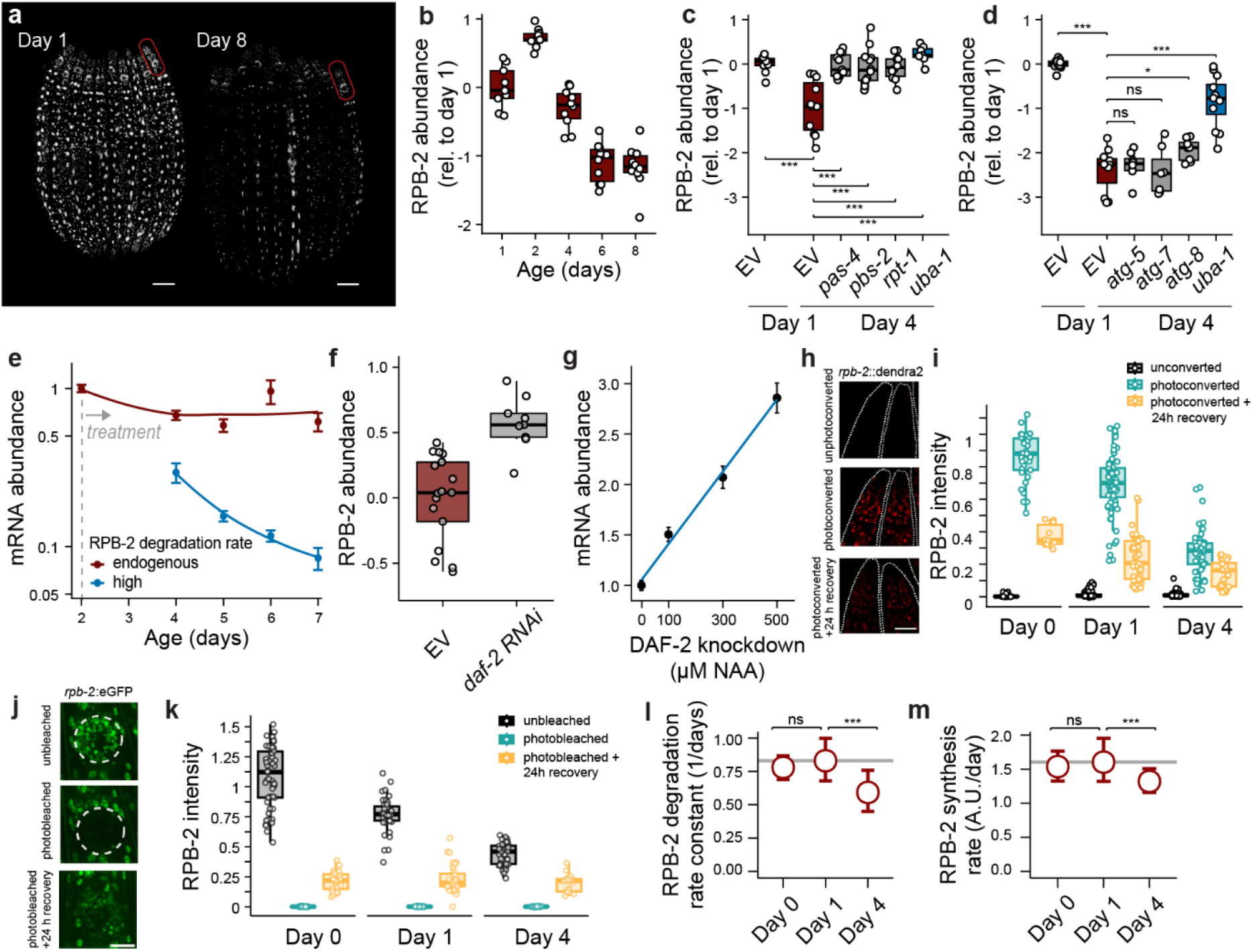
RNA Polymerase II (Pol II) decreases in abundance during aging, driving global decreases in mRNA abundance. **a**. Young (day 1) and old (day 8) *C. elegans* individuals expressing endogenously tagged Pol II subunit RPB-2::GFP, with head regions (*representative region in red)* used for quantification, scale bar = 250 μm. **b**. Fold-change (log_2_) in total RPB-2 abundance in the head, relative to day 1. **c**. Fold-change (log_2_) in RPB-2 abundance following RNA-interference (RNAi) knockdown, starting at the final stage of development (L4), of upstream E1 ubiquitin-activating enzyme (*uba-1*), and downstream ubiquitin-mediated 26S proteasomal and **d**. autophagy degradation components. **e**. Changes in mRNA abundance following *in vivo* acceleration of RPB-2 degradation via an auxin-inducible degron tag relative to young (day 2) individuals. **f**. Changes in whole-body RPB-2 abundance in day 4 populations, following four days of lifespan-extending RNAi knockdown of the insulin/IGF receptor *daf-2* relative to EV control. **g**. Changes in mRNA abundance on day 7 of adulthood, 5 days following NAA treatment for graded increases of DAF-2 knockdown, relative to untreated age-matched individuals. **h**. RPB-2::Dendra2 expression in head regions before *(top)*, immediately after photoconversion *(middle)*, and after 24 hours of recovery *(bottom)*; scale bar = 20 μm. **i**. Quantification of total RPB-2 intensity of the population in *h*. across four biological replicates relative to day 0 (L4) mean. **j**. RPB-2::GFP expression in the posterior pharyngeal bulb before *(top)*, immediately after photobleaching (FRAP) *(middle)*, and after 24 hours of recovery (bottom); scale bar = 20 μm. **k**. Quantification of total RPB-2 intensity of the population in *j*. across two biological replicates relative to the day 0 mean. **l**. The *in vivo* RPB-2 degradation rate constant estimated from photoconversion experiments. **m**. The *in vivo* RPB-2 synthesis rate estimated from photobleaching experiments. For imaging panels *b-d,f,i,k*, each point represents one individual; for mRNA abundance panels *e,g*, each point is the mean of 4 replicates of n=30 each. *** = *P* value <0.001; * = *P* value <0.01.

Outside the context of DNA damage and Pol II stalling^14,15^, the pathways for Pol II degradation are poorly understood. To identify the pathways responsible for Pol II degradation in aging, we tested the ubiquitin-mediated proteasome and autophagy pathways. The age-associated declines in RPB-2 are completely prevented by RNA-interference (RNAi) knockdown of the E1 ubiquitin-activating enzyme *uba-1* (Fig. 2c,d, Fig. S2d) in young adulthood, and also by knockdown of the 26S proteasomal subunits *pas-4, pbs-2*, and *rpt-1* (Fig. 2c). In contrast, knockdown of genes required for autophagy had no effect on age-associated RPB-2 declines (Fig. 2d), indicating that basal Pol II degradation in aging occurs via ubiquitin-mediated 26S proteasome degradation and not autophagy pathways. In total, our results identify a marked decline in Pol II abundance during aging resulting from active, ubiquitin-mediated proteasomal degradation.

To determine whether proteasomal degradation of Pol II is sufficient to drive age-associated declines in mRNA abundance, we used the auxin-inducible degron system to quantitatively increase the ubiquitination rates of RPB-2 over its natural baseline via an orthogonal transgenic E3 ubiquitin TIR1 ligase^16,17^. Increasing RPB-2 degradation rates starting in youth (day 2) accelerated the age-dependent declines in organismal mRNA abundance (Fig. 2e, Fig. S2c), demonstrating that proteasome-mediated degradation of Pol II is sufficient to produce the declines in mRNA abundance observed in aging. Together, our results suggest that age-associated declines in global mRNA abundance arise from a decrease in total mRNA synthesis capacity during aging, a state that can be replicated in young animals by increasing the degradation rate of Pol II.

Because changes in mRNA abundance are tightly coupled to changes in lifespan across interventions (Fig. 1e), we next investigated how perturbations to insulin/IGF signaling affect Pol II abundance. We used RNAi to knock down the insulin/IGF receptor *daf-2*^18^ starting at the end of development (L4 stage), and found that RPB-2 abundance increased with age relative to untreated individuals (Fig. 2f, S2e,f). To measure the influence of insulin/IGF signaling on mRNA synthesis, we knocked down DAF-2 in mid-life starting on day 5, and found that this slowed the rate of age-associated decline in mRNA abundance but did not reverse it (Fig. S2g). Using auxin-inducible depletion, we find that the influence of DAF-2 activity on mRNA abundance is strongly dose-dependent (Fig. 2g, Fig. S2h), highlighting the responsiveness of cellular mRNA abundance to insulin/IGF signaling activity. In total, our data reveal a tight coupling between the activity of the insulin/IGF signaling pathway, the cellular abundance of Pol II, age-dependent declines in mRNA abundance, and organismal aging and lifespan.

### RNA Polymerase II abundance is far above its equilibrium set point at the start of adulthood

The abundance of any protein is determined by the balance between protein synthesis and degradation, which can be studied using the well-established dynamical model dR/dt = K_s_ – K_d_R, in which the abundance of a protein R is increased by synthesis K_s_ and decreased by degradation, K_d_R^19^. For Pol II abundance to decrease over time, the degradation rate must therefore, on average, exceed the synthesis rate during adulthood. To dissect the dynamics of these components, we performed *in vivo* time-series measurements of both the RPB-2 degradation rate using a photoconvertible *Dendra2* tag^20^ (Fig. 2h,i), and the RPB-2 synthesis rate using fluorescence recovery after photobleaching (FRAP)^21^ (Fig. 2j,k) (*See Methods, Suppl. Note 2*). We performed analysis specifically in head regions to minimize known confounding effects of age-related autofluorescence^22^. Our experiments reveal, unexpectedly, that both the RPB-2 degradation rate constant and the RPB-2 synthesis rate significantly decrease with age (Fig. 2l,m)—changes that would be expected to decelerate and accelerate, respectively, age-dependent RPB-2 declines.

To understand how interactions between synthesis and degradation drive losses of RPB-2 during aging, we compared the abundances we measured to the RPB-2 *equilibrium set point*—the steady-state abundance at which RPB-2 protein synthesis and degradation rates are balanced (Fig. 3a). This balance occurs at only a single abundance of RPB-2, defined as R = K_s_/K_d_ whose value can be estimated using experimental data on K_s_ and K_d_. We find that the RPB-2 equilibrium set point remains remarkably constant over the first four days of adulthood (Fig. 3b, Fig. S3a), a period during which total RPB-2 abundance steadily declines about four-fold. These observations indicate that the dynamics of Pol II abundance are driven by a non-equilibrium state of Pol II abundance in youth (Fig. 3c, *Model #1*), rather than by a change in its equilibrium set point (Fig. 3d, *Model #2*). Pol II disequilibrium is a driver of Pol II abundance decline in aging (Fig. 3c), as all individuals exit development with a cellular RPB-2 abundance that far exceeds the steady-state abundance supported by the adult rates of RPB-2 synthesis and degradation. This finding is in striking opposition to the prevailing view that aging involves a departure from homeostatic equilibrium. Instead, our data indicate that young individuals begin adulthood far from RPB-2 equilibrium, but approach the equilibrium set point during aging (Fig. 3e, Fig. S3b).

**Figure 3.**
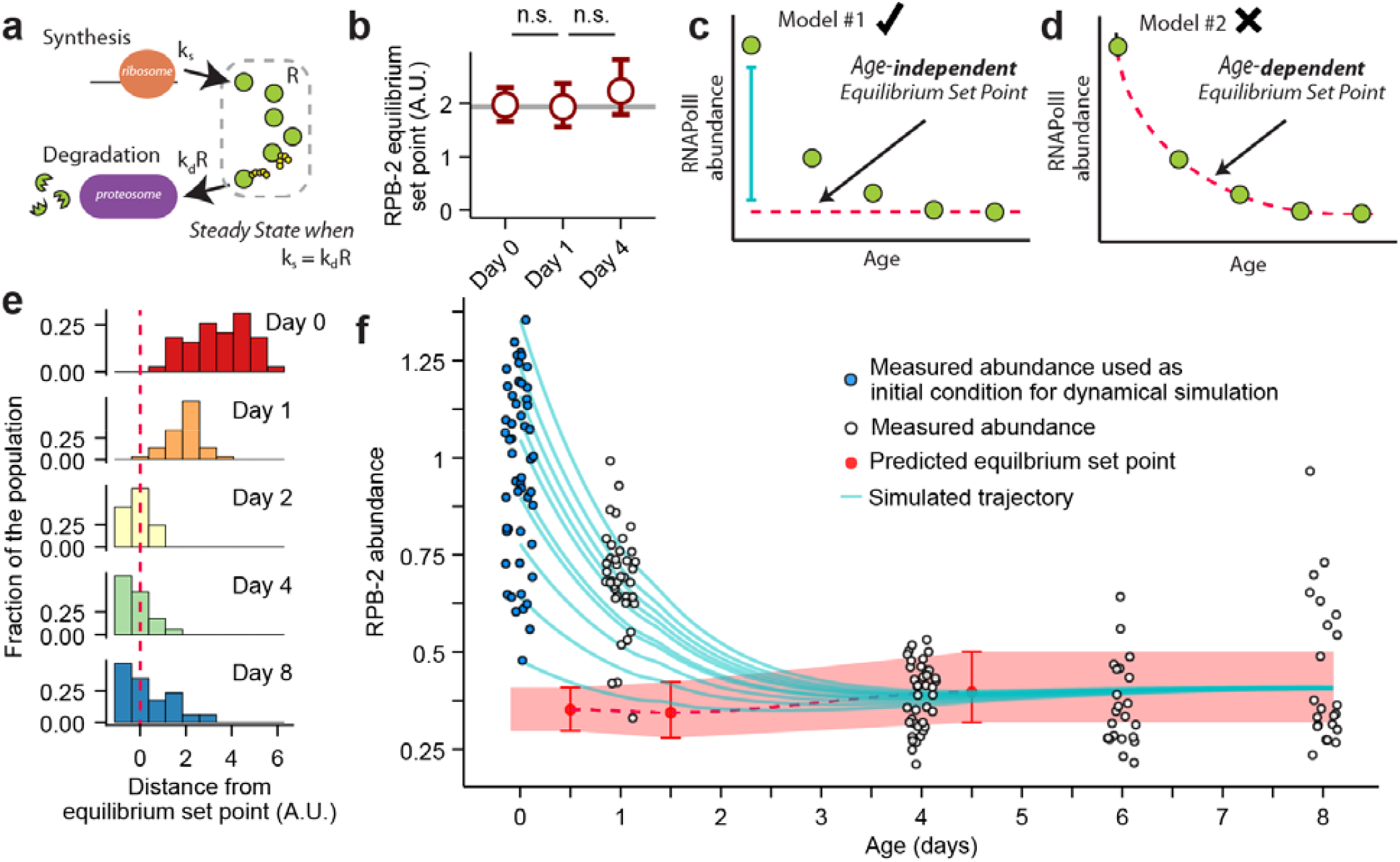
Pol II abundance relaxes towards equilibrium during aging. **a**. The cellular abundance of Pol II subunits is determined by its rate of protein synthesis and degradation, whose balance establishes an equilibrium homeostatic set point for steady-state protein abundance. **b**. Estimates of the *in vivo* RPB-2 equilibrium set point across the first four days of adulthood. **c**. If the equilibrium set point is fixed during adulthood, then any change in RPB-2 abundance *(green circles*) must be driven by out-of-equilibrium phenomena away from steady state *(dotted line)*, and not **d**. by age-dependent changes in the equilibrium set point. **e**. Histograms showing how far each individual’s RPB-2 abundance is from the population equilibrium set point *(dotted line)*, represented as the ratio between individual RPB-2 abundances and the population equilibrium set point. **f**. Comparison of measured RPB-2 abundances *(blue/white circles; each one individual)* to the predicted equilibrium set point *(red points, dotted line)* shown with 95% confidence intervals. To predict the future abundance of RPB-2, measurements at the start of youth (day 0; *blue circles)* were used as initial conditions for a dynamical simulation using protein synthesis and degradation rates. The simulation supports model #1 and predicts exponential declines in RPB-2 abundance *(blue lines)* towards the equilibrium set point, which overlap with measured RPB-2 abundances not used in the simulation *(white circles)*.

One prediction of the disequilibrium model is that young individuals, far from equilibrium for RPB-2, will follow a characteristic relaxation trajectory of RPB-2 abundance towards its equilibrium set point. We confirmed this behavior in a dynamical simulation: using population-average K_s_ and K_d_ estimates in youth, we can predict each individual’s future RPB-2 aging trajectory, which should follow characteristic exponential decays to steady state starting on day 0 (Fig. 3f, *blue curves*). These predicted trajectories match the RPB-2 abundances of peers measured on days 1, 4, 6, and 8 of adulthood (Fig. 3f). If we relax the assumption that all individuals share identical K_s_ and K_d_, and instead consider that RPB-2 abundances in youth result from time-independent heterogeneity in K_s_ or K_d_, our model additionally predicts the inter-individual variance in RPB-2 abundances observed during aging (Fig. S3c,d). In summary, our modelling demonstrates that measurements of the chemical kinetic parameters K_s_ and K_d_ on day 0 are sufficient to predict life-long changes in RPB-2 abundance—strong empirical support for a disequilibrium model (Fig. 3c) in which decreases in RPB-2 abundance during aging are driven by a dynamical relaxation to steady state. In other words, the core components of gene-regulatory machinery required for physiologic homeostasis are themselves out of steady state at the beginning of adulthood.

### The equilibration of RNA Polymerase II abundance is sufficient to drive aging

Finally, we sought to determine the physiological relevance of Pol II dynamics for organismal aging and lifespan. We tested the effect of transient increases in the endogenous degradation rate of RPB-2 at the onset of adulthood (Fig. 4a), hypothesizing that because RPB-2 abundance is far from equilibrium, it would not be subject to a strong homeostatic restoring force capable of recovering from brief losses in abundance. Consistent with this prediction, even transient increases in the ubiquitination rate of RPB-2 starting in young adults produced permanent decreases in mRNA abundance, healthspan, and lifespan in proportion to the duration of increase (Fig. 4b-d, S4e,f). The duration of high RPB-2 degradation was tightly, quantitatively coupled to the magnitude of the effects on global mRNA abundance and lifespan (Fig. 4e), highlighting the crucial influence of Pol II abundance on both transcriptomic and organismal aging.

**Figure 4.**
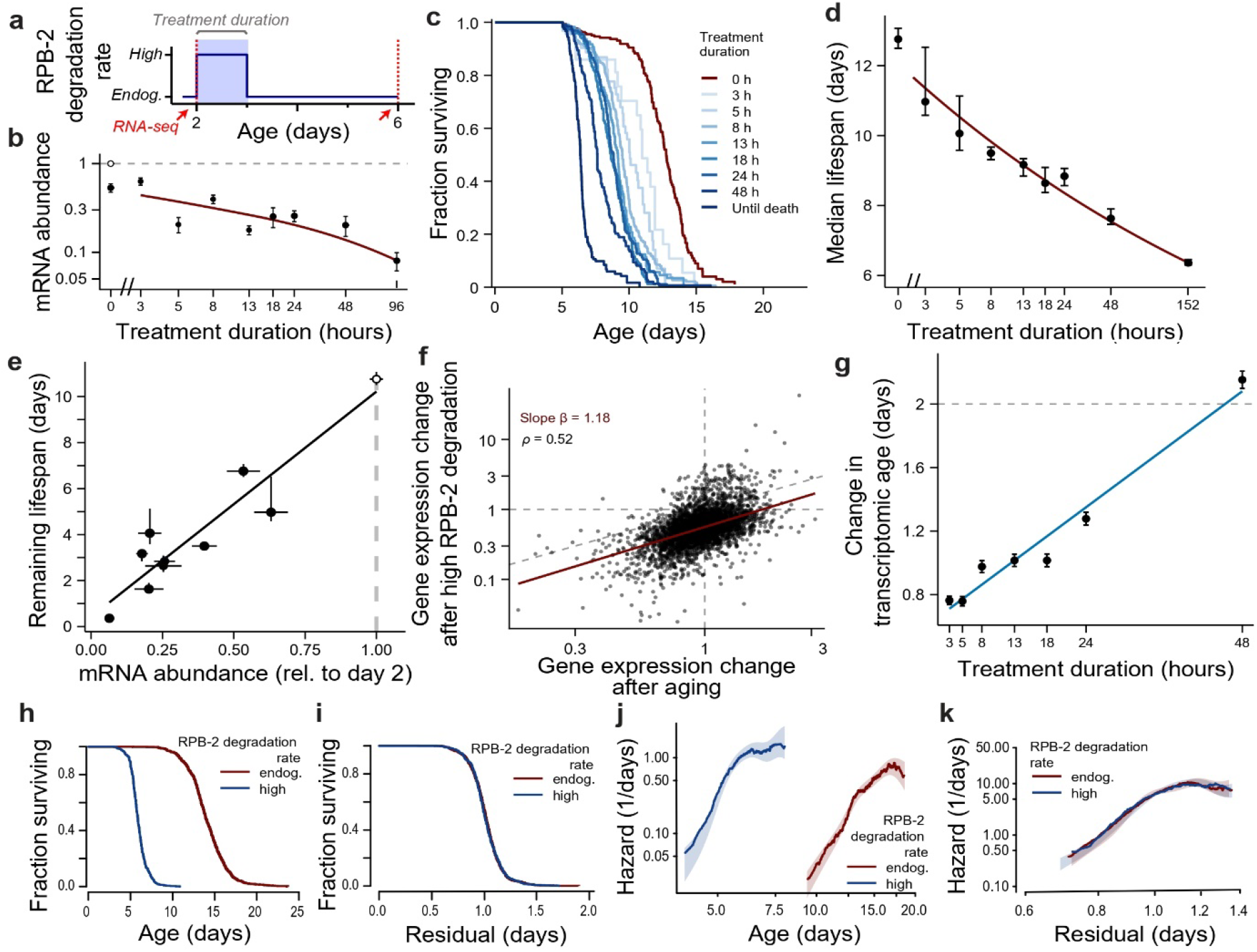
Transient increases in Pol II degradation rates permanently limit global mRNA abundance and lifespan via accelerated aging. **a**. Schematic of experimental design, in which the effects of transient, *in vivo* increases in RPB-2 degradation above the endogenous baseline rate are measured on gene expression (*red arrow; red dotted line*) and lifespan. **b**. The mRNA abundance of populations after transient increases in RPB-2 degradation rates starting on day 2, measured in mid-life (day 6) *(black points)* relative to the young (day 2) untreated population *(white point on dashed line)*, fit with a flexible spline; each point is the mean of 4 replicates (n=30 each). **c**. Kaplan-Meier survival curves showing the effect of transient increases in RPB-2 degradation rates (n=1668). **d**. The median lifespan of the same populations as *b-c*, fit with a non-linear regression curve. **e**. The relationship between mRNA abundance (relative to day 2 untreated animals, *white point; dashed line*) and lifespan in the same populations, with a linear regression line. **f**. The effect of aging. the fold-change in each gene’s expression *(points)* during aging in unperturbed populations (from day 4 to day 6; x-axis). is compared to the effect of RPB-2 degradation (for 48 hours starting on day 2; y-axis) relative to an unperturbed population of the same age (day 4); robust linear fit *(red line)* defines a slope β that captures the relative magnitude of RPB-2’s effect to that of aging. **g**. Using slopes β, we estimate the equivalent number of days of excess aging as a result of high RPB-2 degradation; each point is the mean of 4 replicates (n=30 each) fit with a linear regression line. **h**. Kaplan-Meier analysis comparing the effect on lifespan of life-long, high RPB-2 degradation starting on day 2 of youth (*blue*, n=636) compared to unperturbed populations (*red*, n=1061). **i**. The residuals of an accelerated-failure time (AFT) model fit to the data in *h*., highlighting any deviations from temporal scaling (no significant deviation, *P*>0.2). **j**. Trajectories in the risk of death for populations in *h*. **k**. The same trajectories in the risk of death for populations in *h*., after accounting for differences in timescale using an AFT regression model (no significance, *P*>0.2).

Aging involves not only global declines in mRNA abundance, but also changes in the functional relative expression of individual genes. To determine whether reduced mRNA synthesis accelerates transcriptomic aging, we compared the effect of high RPB-2 degradation rates in young adults to the effects of aging in unperturbed individuals (Fig. 4f). Changes in gene expression induced by high RPB-2 degradation phenocopy the changes produced by aging alone. The relationship between physiologic aging and accelerated RPB-2 losses was also seen in populations exposed to shorter treatment durations, with the magnitude of transcriptomic aging showing a tight dose-dependent relationship to the duration of high RPB-2 degradation (Fig. 4g).

The effects of biological aging can be measured at the demographic level, where changes in the biological aging rate manifest as an acceleration of a population’s age-dependent increase in the risk of death^23,24^. We find that life-long increases in the RPB-2 degradation rate produce a near-perfect temporal scaling^25^ of both survival and hazard-rate trajectories (Fig. 4h-k, Fig. S4a-d), demonstrating that reductions in cellular Pol II abundance accelerate the demographic aging rate. This effect was not limited to lifespan: life-long, high RPB-2 degradation similarly produced a temporal scaling of healthspan (Fig. S4g-j)—indicating that declines in Pol II have systemic effects across multiple outcomes of aging.

## DISCUSSION

Here, we demonstrate that an RNA Polymerase II disequilibrium state at the start of adulthood is sufficient to drive aging. Combining absolute mRNA quantification methods with *in vivo* measurements of Pol II abundance dynamics, we discover a quantitative, causal link between an out-of-equilibrium state of mRNA synthesis capacity present at the onset of adulthood and the progressive, global declines in cellular mRNA abundance that limit lifespan. In *C. elegans*, we demonstrate how changes in mRNA abundance are driven by decreases in mRNA synthesis involving quantitative reductions in the nuclear dosage of Pol II.

Our findings identify preservation of mRNA synthesis capacity as a potential strategy for delaying aging. Interventions that slow declines in Pol II abundance, such as reduced insulin/IGF-1 signaling, preserve global mRNA abundance and extend lifespan, whereas experimentally accelerating age-dependent Pol II loss produces the opposite effect. The coupling between mRNA synthesis capacity, healthspan, and lifespan suggests a new therapeutic goal: stabilizing the disequilibrium that drives aging.

Individuals who exit development out of equilibrium face severe consequences during adulthood. Far from equilibrium in youth, Pol II abundance is not subject to a homeostatic restoring force that would allow individuals to recover from transient perturbations to Pol II. This consequently produces a ratchet-like phenomenon in which declines in Pol II abundance accumulate over time and are not restored, thus accelerating transcriptomic changes, increasing the organismal aging rate, and limiting lifespan. In principle, any molecular state above its equilibrium set point at the start of adulthood represents a physiologic vulnerability—creating a pathological homeostatic process that drives individuals away from youth. Far above steady state there is an asymmetry between damage and repair, in which repair processes that push the state away from the pathological equilibrium set point are opposed by the equilibration kinetics, whereas damage processes that push the state towards pathological equilibrium are compounded.

Pol II is unlikely to be the only process out of equilibrium in youth. First, mRNA synthesis rates are coupled to many other cellular functions, any of which might inherit instability in youth via interactions with Pol II. Second, because development and adulthood place differing physiologic demands on cells, most cellular functions presumably will be quantitatively adjusted at the transition to adulthood. If this adjustment is not instantaneous, then individuals will enter adulthood in a transient state whose equilibration could have pathological consequences. Our results highlight a means for identifying causes of aging based on chemical kinetic disequilibrium states in youth—a quantitative approach for exploring the dynamical origins of age-associated changes in other essential cellular functions.

## ACKNOWLEDGEMENTS

We acknowledge support of the Spanish Ministry of Science and Innovation through the Centro de Excelencia Severo Ochoa (CEX2020-001049-S, MCIN/AEI /10.13039/501100011033) and the Generalitat de Catalunya through the CERCA programme. We thank the CRG Core Technologies Program, including the CRG Advanced Light Microscopy Unit (ALMU), for their support in this work. We acknowledge support from the MEIC Excelencia awards BFU2017-88615-P, PID2020-115189GB-I00, and PID2020-115439GB-I00, and support from the European Research Council (ERC) under the European Union’s Horizon 2020 research and innovation programme (Grant agreement No 852201), and support from the BBVA Programa de Investigación Fundamentos grant CAUSALAGING. Research for this publication has been partially carried out in the Barcelona Collaboratorium for Modelling and Predictive Biology. EMN and this project received funding from the project PID2024-157315NB-I00; the European Union’s Horizon Europe through the European Research Council under the grant agreement number 101042103 and 101187456. OB and ADCF received a CRG Postdoctoral Fellowship to cover the nanopore sequencing costs included in this work. We thank all lab members and Javier Apfeld for helpful feedback on the manuscript.

## AUTHOR CONTRIBUTIONS

ADCF, JV, LS, ME, NO and OB performed experiments. ADCF, JV, KS, LS, NS, and RA performed image analysis. ADCF, KS, NS, OB, and RA performed statistical analysis and modeling. EMN supervised the nanopore sequencing analysis and interpretation. ADCF and NS wrote the manuscript. NS supervised the project.

## SUPPLEMENTARY INFORMATION

Supplementary information for this manuscript includes the following:

- RESOURCE AVAILABILITY
  ∘ Lead contact
  ∘ Materials availability
  ∘ Data and code availability
- MATERIALS AND METHODS
- SUPPLEMENTARY FIGURES S1-S4
- SUPPLEMENTARY NOTES 1-2

