## Supplementary Figures, Methods, and Notes for "Declines in mRNA synthesis set the rate of organismal aging"

### Supplementary Information

This document contains:

- Resource Availability
- Materials and Methods
- Supplementary Figures 1-4
- Supplementary Notes 1-2

#### RESOURCE AVAILABILITY

##### Lead contact

##### Materials availability

Worm strains generated in this study are available from the lead contact upon request.

##### Data and code availability

The short-read mRNA sequencing original data in this study were deposited into the NCBI BioProject database under accession number PRJNA1474372. The long-read mRNA sequencing data (basecalled FASTQ files), as well as processed per-gene counts, have been deposited into the Gene Expression Omnibus (GEO), under accession code GSE334229. The code is available on GitHub, https://github.com/nstroustrup/mRNA_abundance

#### MATERIALS AND METHODS

**Nematode maintenance**—*C. elegans* were maintained on Nematode Growth Medium (NGM) plates seeded with *E. coli* NEC937 (OP50 ΔuvrA; KanR) and kept at 20°C, except for sterile populations which were transferred to 25°C for germline ablation induction by temperature-sensitive mutation *glp-1(e2141)*. UV inactivation of NEC937 was achieved by irradiating freshly grown liquid bacteria with 100 μJ/cm$^2$ in a spectrophotometer (Ultraspec 3100 Pro).

**Lifespan and healthspan assays**—Survival assays were performed using the Lifespan Machine[1] at a population density of 40 to 60 animals per plate. Age synchronous populations for non-imaging experiments were obtained by hypochlorite treatment of day 2 gravid animals. For imaging experiments, populations were synchronized through a lay-off. For non-sterile populations, animals were transferred at the late L4 stage to agar plates containing 10 μg/ml 5-fluoro-2'-deoxyuridine (FUDR, Sigma) to eliminate live progeny.

For auxin-inducible degradation[2] experiments, auxin-analogue 1-Naphthaleneacetic acid (NAA; Sigma-Aldrich) was solubilized in 1M potassium hydroxide and added to molten agar at a final concentration of 375 µM for RPB-2 degradation and 500 µM for DAF-2 degradation with treatment starting on day 2 of adulthood, unless otherwise stated. For all RNA interference (RNAi) experiments, treatment started on the last development stage (L4). All imaging and transcriptomics nematode populations consider germline-ablated *glp-1(e2141)* individuals unless otherwise stated.

**Nematode strains**—The following *C. elegans* strains were used: QZ0 (Bristol N2), CB4037 (*glp-1*(*e2141*) III), CF1880 (*glp-1*(*e2141*) III; daf-16(*mu86*) I), CER510 (*rpb-2*::GFP(dpiRNA)::AID::3xFLAG III), AMP306 (*rpb-2*::eGFP::AID::3xFLAG(*ohm50*) III; *glp-1*(*e2141*) III), AMP100 (ieSi57[*eft-3p*::TIR1::mRuby::*unc-54* 3'UTR + *Cbr-unc-119*(+)] II; *rpb-2(ohm20)*[*rpb-2*::AID::3xFLAG(codon optimized)] III), AMP159 (ieSi57[*eft-3p*::TIR1::mRuby::*unc-54* 3'UTR + *Cbr-unc-119*(+)] II; *rpb-2(ohm20)*[*rpb-2*::AID::3xFLAG(codon optimized)] III; *glp-1(e2141)* III), AMP254 (*rpb-2)*[*rpb-2*::eGFP::AID::*3xFLAG(ohm50)*] III; *glp-1(e2141)* III; ieSi57[*eft-3p*::TIR1::mRuby::*unc-54* 3'UTR + *Cbr-unc-119*(+)] II), AMP116 (weSi174[*Peif-3.B*::TIR-1::linker::mCherry(dpiRNA)::*tbb-2* 3'UTR; *unc-119*(+)] II; *daf-2(syb1177)* [*daf-2*::AID::TEV::3xFLAG] III; *glp-1(e2141)* III), AMP340 (*rpb-2::*Dendra2(*ohm80*) III; *glp-1(e2141)* III).

**Short-read mRNA sequencing—**Single-individual and bulk samples were processed following published methods[3]. mRNA sequencing was performed using the Smart-Seq2 protocol[4]. Lysis buffer was prepared with ERCC spike-ins[5] added to a final dilution of 1:40,000 for both single-individual and bulk sequencing. For bulk sequencing, 30 synchronized animals were picked into 120 µL lysis buffer, with four replicates each. For single-individual sequencing, individual animals were picked into 80 µL lysis buffer. Immediately after collection, nematode suspensions were shock-frozen into liquid nitrogen and stored at -80°C. Lysis was performed at 65°C for 10 minutes, followed by 85ºC for 5 minutes for enzyme deactivation. Complementary DNA (cDNA) libraries were generated using the Smart-Seq2 protocol. Purification was conducted with in-house SPRI-style paramagnetic beads designed to function similarly to AMPure XP beads (Beckman Coulter), using a bead-to-sample ratio of 0.8. The resulting libraries were analyzed for fragment size distribution using the TapeStation 4150 system (Agilent), and cDNA concentrations were quantified with the Quant-it kit (Invitrogen) on a plate reader (Tecan). The Nextera DNA library prep protocol (Illumina) was used to generate Nextera sequencing libraries, using tagmentation of the cDNA and PCR amplification with indexing primers. Libraries were then purified twice using the in-house SPRI paramagnetic beads with a 0.9 bead-to-sample ratio. The resulting fragment size distribution was re-analyzed with TapeStation 4150 (Agilent), and library concentration was measured again using Quant-it (Invitrogen) on a plate reader (Tecan). The resulting RNA-seq libraries were pooled to equal mass, and the paired-end Nextera libraries were sequenced using Illumina NextSeq500 using high-output 75-cycle v2.5 kits (Illumina).

RNA-seq reads were aligned using STAR version 2.6.0c[6] to the *C. elegans* WormBase reference genome (WS265), with modifications to include ERCC spike-in sequences. Gene count quantification was performed using featureCounts version 2.0.0[7]. Genome coverage of reads was quantified with BEDTools[8].

**Long-read direct mRNA sequencing** –Total RNA was extracted using TRIzol LS reagent (Thermo Fisher #10296028) followed by purification with the Zymo Research RNA Clean & Concentrator Kit (Zymo, #R1013). Nematode samples in 330 µl M9 collection buffer supplemented with magnesium and ERCC spike-ins[5] (0.5%) were mixed with 1 mL TRIzol LS, flash-frozen by immersing the tube bottom-first into liquid nitrogen, and thawed in a 37ºC heat-block. This freeze–thaw cycle was repeated seven times. Following the final thaw, samples were subjected to five cycles of vortexing (3200 rpm, 30 seconds on/30 seconds off). Chloroform (267 µl) was added, samples were vortexed for 30 seconds, and centrifuged at maximum speed at 4ºC for 15 minutes. The aqueous phase was transferred into a new tube and re-extracted with an equal volume of chloroform, then vortexed and centrifuged under the same conditions, to ensure complete removal of organic contaminants. The final aqueous phase was mixed with an equal volume of absolute ethanol and applied to Zymo-Spin columns for cleanup following the manufacturer's instructions.

Direct RNA libraries were prepared using the Oxford Nanopore Technologies Direct RNA Sequencing Kit (SQK-RNA004) following the manufacturer's protocol with minor modifications. Briefly, 300 ng of total RNA isolated from *glp-1(e2141) C. elegans* collected at days 2, 8 and 13 of adulthood ages was ligated to different RT adapter (RTA) sequences for multiplexing. Ligation was performed using T4 DNA ligase (NEB, #M0202M) at room temperature for 15 min, followed by reverse transcription with SuperScript IV (Invitrogen, #18090050). Barcoded libraries were pooled and purified using RNAClean XP beads (Agencourt, #A63987) and quantified using the Qubit RNA HS Assay (Thermo Fisher Scientific, #Q32855). The pooled library was then ligated to the RNA ligation adaptor (RLA), cleaned up using RNAClean XP beads, and loaded onto PromethION flow cells (RNA004 chemistry) for sequencing (up to 72 h).

***in vivo* imaging:** Nematodes were anesthetized on 5% agarose pads using 5 mM levamisole (ITW Reagents, P110801) dissolved in M9 and secured with a #1.5 coverslip.

**Fluorescence Recovery after Photobleaching (FRAP) and Dendra2 imaging**—Measurements of synthesis rates in *rpb-2*::eGFP;*glp-1(e2141)* were achieved by FRAP, where images of the nuclei-dense region in the entire posterior bulb of the pharynx were collected immediately before and after exposure to 100% of 488 nm laser for 1 second using a UGA-42 Firefly Point Scanning Device (Rapp OptoElectronic). Measurements of degradation rates in *rpb-2*::Dendra2;*glp-1(e2141)* strains generated in this study, were achieved by utilizing the photoconvertible fluorophore Dendra2[9], where images of the head regions of nematodes (from the most anterior part until the posterior bulb of the pharynx) were collected immediately before and after photoconversion of Dendra2 by exposure to one minute of blue light illumination using a CoolLED pE-300 Ultra system with a DAPI filter. Individuals recovered immediately after imaging were washed three times with M9 and recovered into freshly seeded plates for re-imaging 24 hours later.

Images were acquired using the Evident IXplore IX83 SpinSR Super Resolution Microscope System equipped with a UPLSAPO30xS 1.05 NA silicon oil immersion objective and 488 nm and 561 nm Coherent diode lasers controlled by the OLYMPUS cellSens Dimension imaging software (version 4.3). Images were captured with the 488 laser power set at 30% and 100 ms of exposure time, while the 561 laser power was set at 70% and 200 ms of exposure time. Tiled Z-stacks were captured using a step size of 1 μm. The 488 nm laser was used to capture the pre-converted (green) RPB-2::Dendra2 signal and to measure remaining signal post-photoconversion. The 561 nm laser was used to capture post-photoconversion (red) of RPB-2::Dendra2 signal as well as to capture autofluorescence on pre-photoconversion animals.

#### Statistical methods for RNA sequencing

**Estimation of total mRNA abundance from short-read sequencing:** The complete derivation, modelling assumptions, and residual-based diagnostics for mRNA abundance quantification are provided in *Suppl. Note 1*. Total mRNA abundance was estimated as the ratio of an mRNA-derived scale factor and an ERCC-derived technical depth factor, both obtained from regression rather than from raw count sums. The mRNA scale factor was estimated by fitting a pooled linear model of log-transformed mRNA counts on sample identity, using all retained genes across all samples in a given experiment (RcppArmadillo::fastLm). The first sample of the youngest (or untreated) condition was used as the reference, so that each sample-specific coefficient represents the multiplicative log-scale displacement of that sample's full mRNA count distribution relative to the reference. Unlike summed mRNA counts, this estimator is insensitive to highly expressed outlier genes; unlike scran size factors or library-size normalization, it does not impose a constant-total constraint, and therefore preserves rather than removes genuine global shifts in mRNA abundance. The ERCC depth factor was estimated sample-by-sample by regressing log spike-in counts on log known input concentrations, the latter entered as an offset. Because all ERCC species within a sample share library preparation, capture, and sequencing conditions, this regression isolates a single per-sample technical scaling parameter while using the full set of spike-ins jointly rather than discarding per-species information by summing. Standard errors from both regressions were propagated to the mRNA abundance estimate via the first-order delta method under approximate numerator–denominator independence, and 95% confidence intervals were reported on the raw scale. mRNA abundance values are shown throughout the figures relative to the mean of the youngest or untreated reference group. Tissue-specific transcripts were identified using a reference set of mRNA collected from FACS-sorted cells[10].

**Estimation of total mRNA abundance from long-read sequencing:** Raw signal data (POD5) from two PromethION flow cells were demultiplexed with SeqTagger (standalone) using the b100 RNA004 model set, and reads assigned to experimental barcodes were retained for downstream analysis using a base quality filter (baseQ ≥ 50). Basecalling was performed using Dorado (v1.0.2) with the RNA004 SUP model (rna004_130bps_sup@v5.2.0), producing barcode-specific read sets.

Reads were aligned using a two-step minimap2 strategy to separate spike-in and rRNA reads from genomic alignments. First, reads were mapped to a combined reference containing *C. elegans* rRNA sequences together with ERCC/SIRV spike-in sequences using ONT-optimized parameters (-ax map-ont). Reads unmapped in this step were subsequently aligned to the *C. elegans* reference genome (WBcel235) using spliced alignment optimized for direct RNA (-ax splice -uf -k14). Alignments were sorted and indexed with samtools, and the rRNA/ERCC/SIRV and genome alignments were merged to generate final per-barcode BAM files for quantification.

Gene-level quantification was performed using featureCounts (Rsubread; long-read mode) with a curated GTF annotation, extracting gene identifiers and associated attributes (gene name and gene biotype). For downstream quantification of total mRNA and ERCC-normalized abundance across aging, ERCC spike-in counts were summed per sample and used to compute nematode-aware ERCC scaling factors (based on the number of nematodes per timepoint). Gene counts were then normalized to ERCC per individual nematode, and total mRNA abundance was summarized relative to day 2 for visualization and comparative analyses.

**Sample and feature filtering**: Sample and gene (feature) filtering thresholds were adjusted to the sequencing depth across experiments, biological complexity (single nematodes, pooled nematodes, mouse tissue pseudo-bulks), and ERCC input ratio. mRNA features were retained if expressed above a per-count threshold in at least 95% of remaining samples, and ERCC species were retained under the same presence criterion. The specific per-panel implementation details can be found on GitHub.

***M. musculus* tissue and single-cell mRNA abundance analysis**: Tissue-level mRNA abundance in mouse was estimated from cell-count-matched pseudo-bulks rather than from individual cells, to avoid confounding by differential cell recovery between 3- and 18-month animals. We focused on tissues for which at least 100 cells had been collected at both ages. To eliminate the influence of differences in cell number across ages, the same number of cells were selected at each age—all ages were selected by sampling without replacement (during bootstrapping) the minimum number of cells present at any age. Then UMI counts were summed across cells to yield one mRNA pseudo-bulk per tissue × age combination, with ERCC counts aggregated over the same cells. Pseudo-bulks were filtered as described above and passed through the mRNA abundance estimator. ERCC species whose mix 1 input concentrations fell outside the linear log–log dynamic range (retained range: 0.5–10,000) were excluded prior to fitting to prevent saturating or near-zero spike-ins from biasing the depth regression.

The same cell-count-matched pseudo-bulk procedure was applied at the level of cell-type. The same cell-number balancing procedure performed for tissues was repeated for individual cell types.

Cells were grouped by their cell ontology. Cells lacking an ontology assignment were excluded. For each cell type with at least 100 cells in each age group, the minimum cell count across age groups was taken as a fixed pool size. This number of cells was sampled without replacement from each age and their UMI counts summed to yield one mRNA pseudo-bulk per cell type × age combination, with ERCC counts aggregated over the same cells. Pseudo-bulk filtering, ERCC range restriction, and mRNA abundance estimation were performed exactly as for the tissue-level analysis.

To confirm that age-associated differences were not confounded by age-independent inter-individual heterogeneity—which is a concern due to the low number of mice considered—we confirmed our mRNA abundance estimates using subsets of cells obtained from single mice at each age (young, old).

***C. elegans* single-cell and single-nuclei mRNA abundance analysis**: In the *C. elegans* single-cell and single-nuclei mRNA datasets, mRNA abundance was estimated from total UMI counts per cell (single-cell) or per nucleus (single-nucleus): within each cell type or tissue, the mean total UMI count was computed at each age, and the ratio of these means between ages was taken as the relative change in mRNA abundance. This rests on the assumption that per-cell capture efficiency is comparable across ages within a given cell type, such that age differences in mean UMI count reflect differences in captured mRNA rather than differences in capture.

For the single-cell data, cells were assigned to curated cell types, and cell types flagged for exclusion or represented by fewer than 15 cells or fewer than 200 detected genes were discarded. For each retained cell-type, the relative mRNA abundance with age was computed as the ratio of mean total UMI counts between ages. A 95% confidence interval was obtained by resampling cells with replacement within each age over 10,000 bootstrap iterations and taking the 2.5th and 97.5th percentiles of the resulting ratio distribution.

For the single-nuclei data, tissues with fewer than 40 nuclei were excluded. mRNA abundance was computed per tissue as the ratio of mean total UMI counts per nucleus between ages, applying 95% confidence intervals from 10,000 bootstrap resamples of nuclei within each age.

**Rate of decline in mRNA abundance correlation with lifespan**: For each of the 13 *C. elegans* genetic and environmental conditions, the rate of decline in mRNA abundance was defined as the negative slope of an ordinary least-squares linear regression of log2(mRNA abundance) against chronological age (t) in days, according to the model log2(mRNA abundance) (t) $= mt + b$, where $m$ represents the rate of change in mRNA abundance per day and $b$ the predicted mRNA abundance at $t = 0$. The resulting decline rates were correlated with remaining lifespan from day 2 using Pearson correlation. All conditions share a common *glp-1(*e2141) genetic background and were grown on UV-inactivated bacteria unless live bacteria is mentioned, both axes are displayed on a log scale.

#### Statistical methods for image analysis

**Fluorescence quantification**—Fluorescence intensity was quantified for each experiment by determining a minimum, common threshold value of each fluorophore color channel to subtract from all pixels, where signal below the threshold was considered noise. All positive number of pixels were summed within the drawn mask of the area to quantify. For body size measurements, masks of the area in whole individual nematodes were drawn and quantified as the sum of all pixels. For experiments where only head regions were imaged, masks were drawn from the most anterior part of the body until the posterior end of the posterior bulb. Final fluorescence intensity or body size area are displayed as the log2 of the sum in arbitrary units (A.U.).

**Estimating absolute RPB-2 abundance**—To estimate the background intensity produced by technical factors and autofluorescence, we identified the 20 by 10-pixel rectangle with the lowest mean brightness in each image. We then applied a linear rescaling to all pixels such that the 10% quantile of the dimmest rectangle is set to the value 300 and the 90% quantile to 700. Because RPB-2::eGFP is measured in the green channel, we used the red channel to determine autofluorescence, which was then removed by setting the green channel value to 0 wherever the red channel exceeded a manually chosen value. Images were captured using the LSM980 inverted microscope (Zeiss) equipped with the Airyscan2 detector (Zeiss) using a dry 10X objective. Images were captured with the 488 laser power set at 50% and 650 V for measuring eGFP, while the 561 laser power was set at 70% and 700 V for measuring autofluorescence. A tiled Z-stack at 1 μm steps was captured using the extended focal range mode.

**Batch correction of RPB-2 abundance following *daf-2* knockdown**—To batch-correct across four independent biological replicates of RPB-2 abundance following *daf-2* RNAi, we fit the following linear model: log(mean expression) = $\beta_0$ + β_{RNAi}(RNAi condition) + β_{batch}(batch) + ε, where $\varepsilon \sim \mathrm{Normal}(0, \sigma^2)$. The RNAi condition term captures the biological effect of *daf-2* knockdown, while the batch term captures replicate-specific technical variation.

**Estimation of RPB-2 synthesis rates, degradation rate constants, and equilibrium set point**—Changes in nuclear RPB-2::eGFP and RPB-2::Dendra2 fluorescence were compared immediately after photoconversion or photobleaching, respectively, to the same population after 24-hours of recovery. Degradation rates were estimated by quantifying the drop in Dendra2 fluorescence in the red channel during the 24-hour recovery period. Synthesis rates were estimated by quantifying the increase in RPB-2::eGFP fluorescence during the 24 hour recovery period.

To account for day-to-day variation in autofluorescence while estimating degradation rates given that the Dendra2 fluorophore has a lower signal-to-noise ratio than other fluorophores used, we used adaptive thresholding for this data. For each individual, we computed the $0.999^{th}$ percentile of brightness. We then used each of these as the threshold value to subtract from the image and summed all positive pixels. The full computational approach for using these changes in fluorescence intensity to estimate the synthesis rate, degradation rate and the degradation rate constant, including bootstrapping to estimate statistical confidence across all technical and biological variation, is described in detail in *Suppl. Note 2*. The population-average equilibrium set points were estimated as the ratio of the synthesis rates and degradation rate constants.

**Dynamical simulation of future RPB-2 abundance trajectories**—The trajectory of individuals governed by the dynamical equation $dR/dt = k_s(t) - k_d(t)R(t)$ was estimated using numerical integration. $k_s(t)$ and $k_d(t)$ were specified as a linear interpolation between the population-average values measured in FRAP and Dendra2 experiments, respectively, at each experimental time-point. The simulation was run N times, once for each of the N individuals in the day 0 population, with $R(t)$ set to that individual's RPB-2 intensity.

In Supplementary Fig. 3c-d, we explored the hypothesis that individual differences at day 0 correspond to proportional, fixed (*i.e* time-independent) differences in individuals' synthesis and degradation rates. For both panels, the distance of individual *i* from the population average on day 0 was calculated as $Z^i = R(0)^i / \left(\frac{1}{N}\sum_i^N R(0)^i\right)$. For Suppl. Fig. 3c, we assumed individuals differ in respect to $k_d$, which then suggests the "individualized" dynamical equation $dR/dt = k_s(t) - \frac{1}{Z^i} k_d(t)R(t)$ . For Suppl Fig. 3d, we assumed individuals differ in respect to $k_s$ which then suggests $dR/dt = Z^i\, k_s(t) - k_d(t)R(t)$ .

### Supplementary Figures

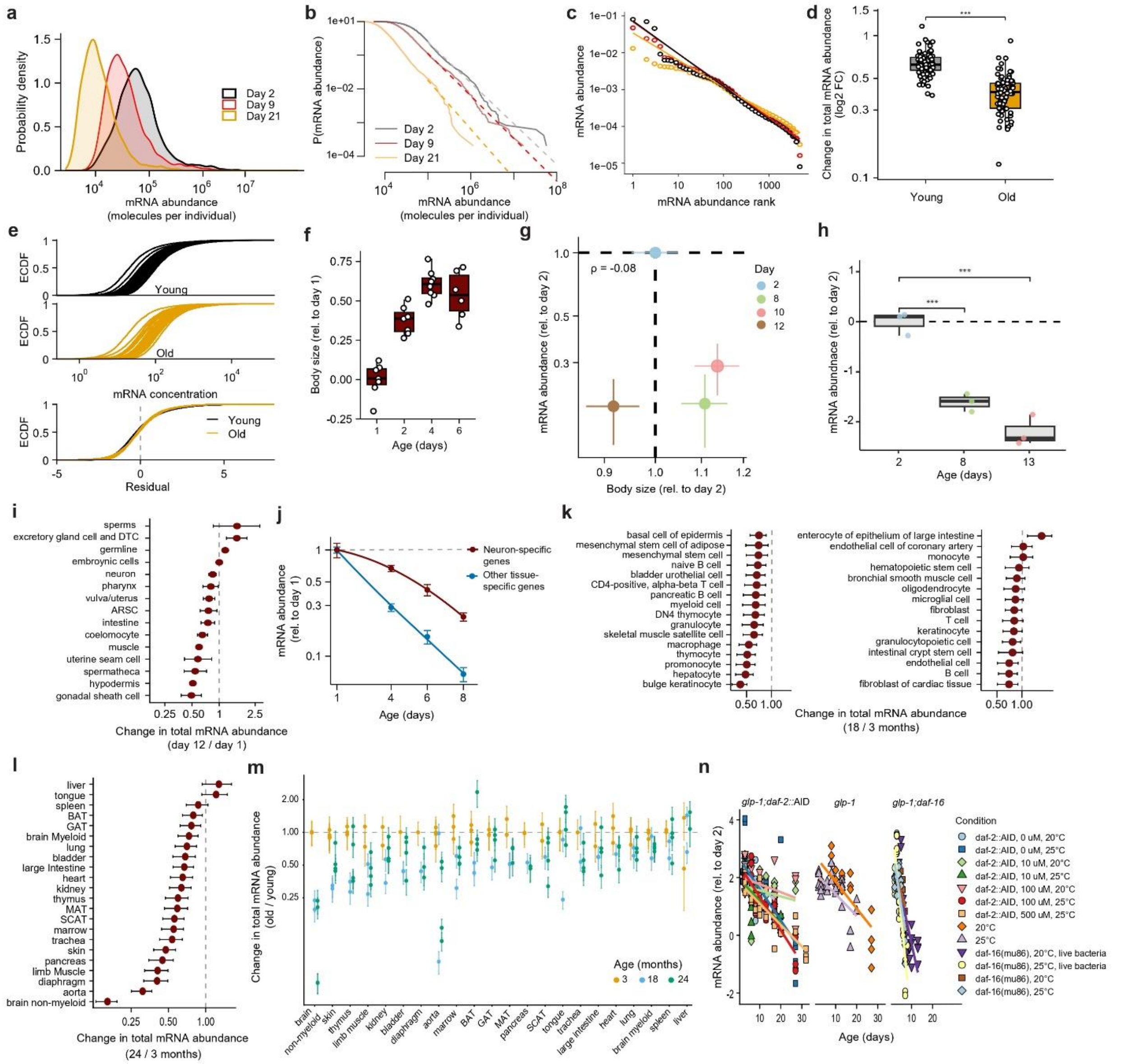


**Figure S1—The mRNA abundance of cells and organisms decreases during aging and the rate of decline closely tracks lifespan. a.** The distribution, across the transcriptome, of the number of copies for each mRNA molecule in an individual wild-type *C. elegans* in young (day 2), old age (day 9) and very old age (day 21), measured as the mean of 4 replicates (n=30 each). **b.** A diagnostic exploring the same mean mRNA abundance distributions in *a*, on transformed axes that linearize power-law distributions, with a linear fit (dotted lines) repeating the analysis of Ueda *et al.* (2004). **c.** A second diagnostic of populations in *a*, that ranks genes in descending order of their absolute mRNA abundances, to identify instances of Zipf's law as described in Furusawa and Kaneko (2003). **d.** The total mRNA abundance measured separately in each *C. elegans* individual (*one point*) in young (day 1) and old (day 8) germline-ablated *glp-1(e2141)* populations. **e**. The same single-individual transcriptomes in *d.*, plotted as cumulative distribution functions of mRNA abundances in young, day 1 *(top)* and old day 8 *(middle)* populations. A log-normal generalized linear regression model was applied to test if these distributions differ only by scaling, in which case model residual abundances *(bottom)* should overlap. *P* value of residuals for age-matched individuals = >0.01; *P* value between young/old individuals = <0.01; modified Kolmogorov-Smirnov (KS) test. **f.** Changes in body size, measured using confocal imaging, in single wild-type *C. elegans* individuals (*each point*), *during aging* relative to young (day 1). **g.** A comparison between changes in body size and mRNA abundance during aging relative to young (day 2) in germline-ablated *glp-1(e2141)* individuals, from the mean across 4 replicates (n=30 each); Pearson correlation = -0.08. **h.** Estimation of total mRNA abundance in *glp-1(e2141)* individuals relative to young (day 2) using long-read nanopore direct mRNA sequencing as an orthogonal measurement (*Methods*). **i.** Change in the mean total mRNA abundance of single nuclei between old (day 12) and young (day 1) wild-type *C. elegans* individuals, obtained by single-nuclei sequencing in Gao *et al*. (2024). **j.** Total abundance of transcripts in *glp-1(e2141)* populations identified as being expressed exclusively in neurons compared to all other tissues, mapped with "tissue-ome" annotations of Kaletsky *et al*., (2018), measured as the mean of 4 replicates (n=30 each). **k.** Data from the same single-cell *M. musculus* experiments of Tabula Muris Senis (2020) shown in Fig.1c, but pseudo-bulking cells by cell-type rather than tissue-of-origin. **l**. Change in mean total mRNA abundance, as in panel *k*, but considering cells isolated from older mice (24 months) relative to young (3 months) wild-type male individuals, grouped by tissue-of-origin. **m**. Change in total mRNA abundance, as in panel *k* and *l*, plotted by individual mice and across ages relative to young adult mice; each point represents one individual. **n**. The estimated slopes in mRNA abundance across all lifespan-altering interventions shown in Fig.1e; each symbol n=30.

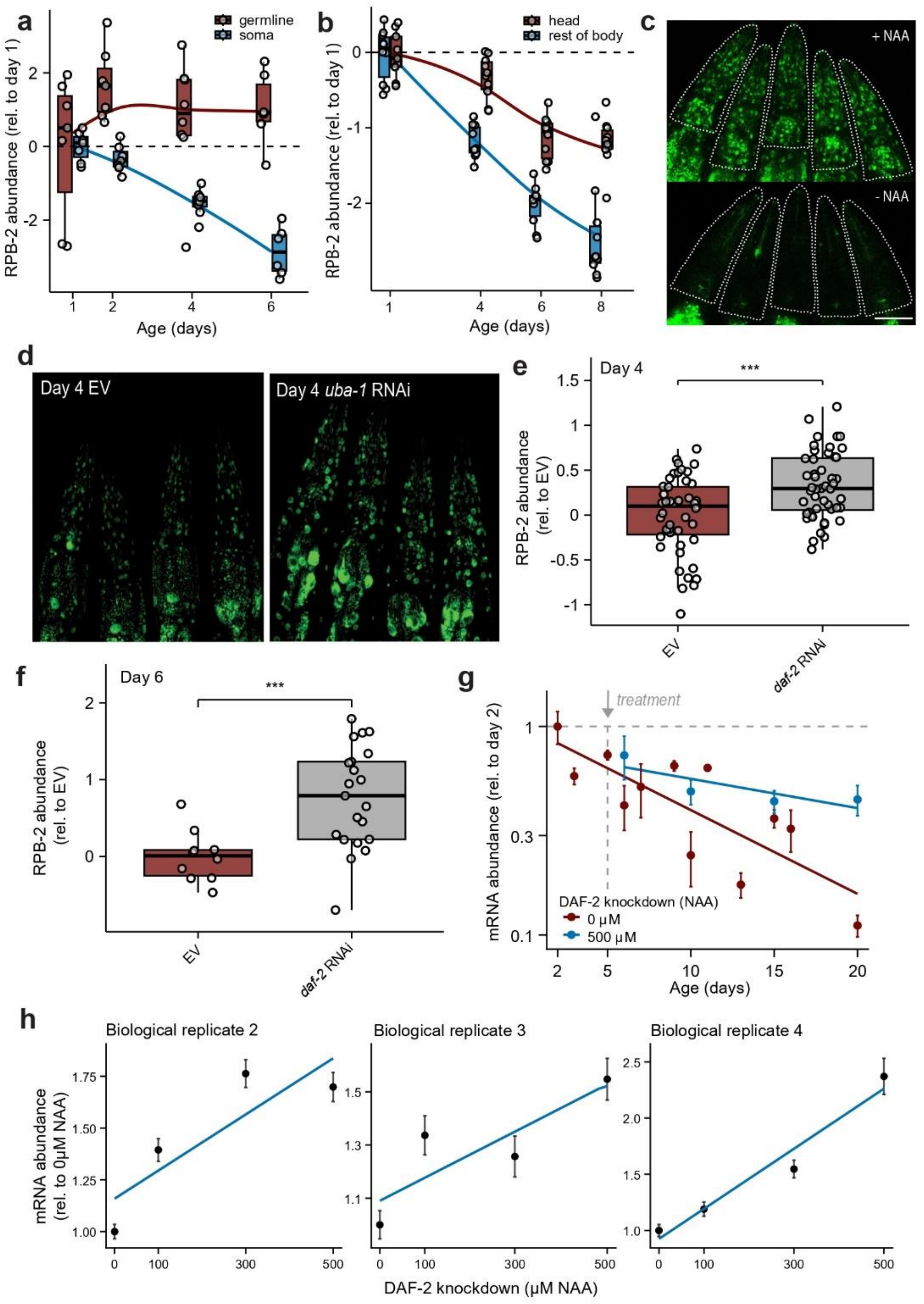


**Figure S2—RNA Polymerase II (RNAPolII) decreases in abundance during aging, driving global decreases in mRNA abundance. a.** Changes in RPB-2 abundance during aging relative to young (day 1) wild-type (N2) *C. elegans*, measured in all somatic nuclei compared to germline nuclei across the two gonad arms (*each point is one individual*). **b.** Changes in RPB-2 abundance during aging in germline-ablated *glp-1(*e2141) individuals (*each point)*, measured in head nuclei only compared to all other somatic nuclei. **c.** The effect of increased RPB-2 degradation for 48 hours on RPB-2 expression (shown in head regions) of day 4 individuals. Residual fluorescence in the foregut is the result of autofluorescence. **d.** The effect on RPB-2 expression of RNA-interference (RNAi) knockdown of the E1 ubiquitin-activating ligase *uba-1* for 4 days starting on L4 (head regions shown)**. e.** Changes in head-nuclei RPB-2 abundance on day 4 populations, following four days of lifespan-extending RNAi knockdown of the insulin/IGF receptor DAF-2, batch-corrected across four independent biological replicates (*each point is one individual*). **f**. Changes in head-nuclei RPB-2 abundance on day 6 populations, following six days of DAF-2 knockdown (*each point is one individual*). **g.** The effect on mRNA abundance of NAA-induced DAF-2 knockdown starting on day 5 compared to untreated animals, measured as the mean across 4 replicates (n=30 each). **h.** Additional biological replicates of the DAF-2 knockdown experiment in Fig.2g, showing the dose-dependent relationship between DAF-2 knockdown and mRNA abundance on day 7, following 5 days of graded DAF-2 knockdown (each point is the mean of 4 replicates, n=30 each). *** = *P* value <0.001.

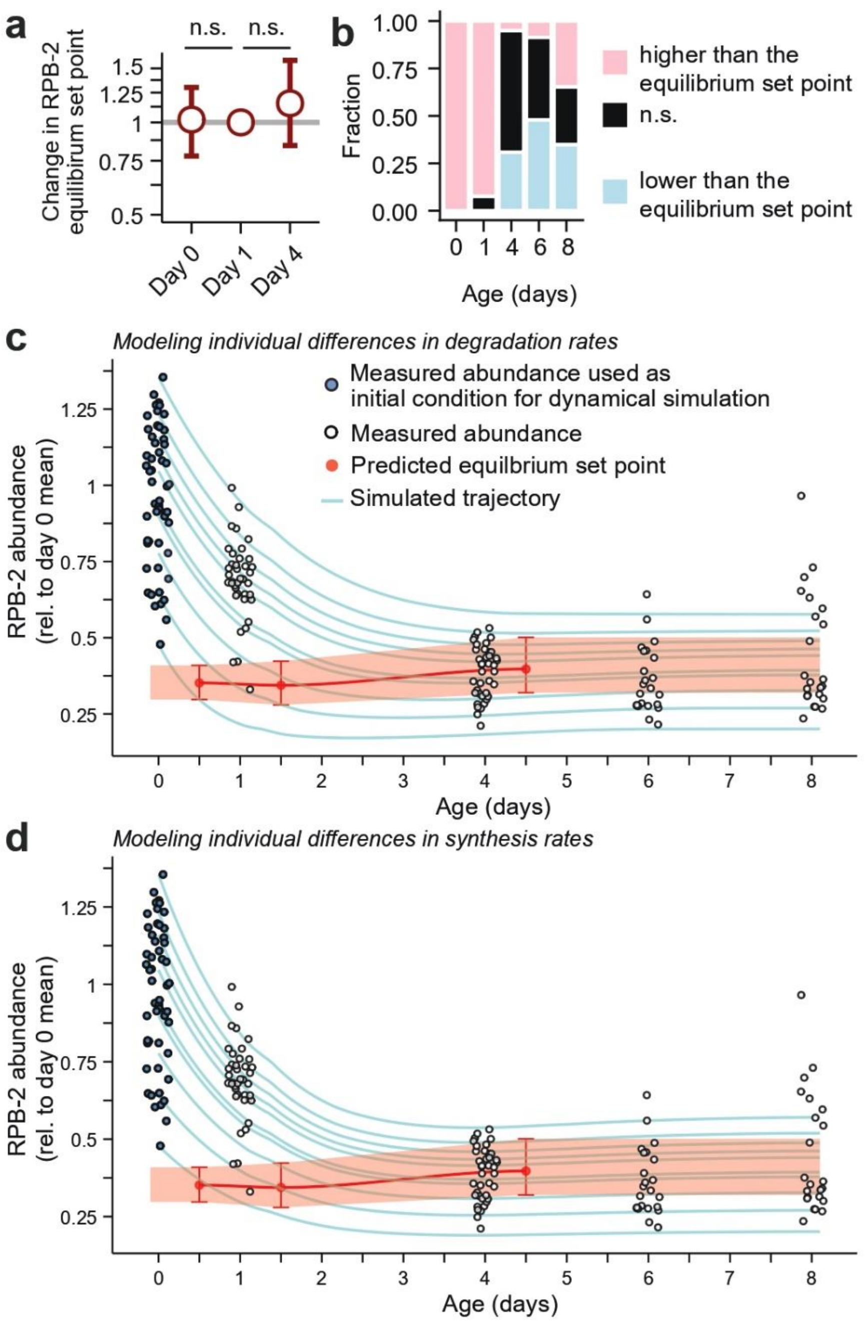


**Figure S3—RNAPolII abundance relaxes towards equilibrium during aging. a.** Estimates of the age-dependent change in the RPB-2 equilibrium set point, with bootstrap 95% confidence intervals. **b.** The fraction of individuals on each day whose RPB-2 abundance is significantly higher *(pink)*, significantly lower *(blue)*, or statistically indistinguishable *(black)* from the population's estimated equilibrium set point. **c**. The same data and equilibrium set point estimates presented in Fig.3f, but with different frailty components. To consider the possibility that inter-individual heterogeneity in RPB-2 abundances observed at the start of youth (day 0) reflects age-independent differences in RPB-2 degradation rate constants ($k_d$), we assigned each individual a unique $k_d$ proportional to the deviation of that individual's RPB-2 abundance from the day 0 population average. **d**. The same data and equilibrium set point estimates as presented in panel *c*, but assuming time-independent heterogeneity in individual RPB-2 synthesis rates ($k_s$), where each individual is assigned a unique $k_s$ proportional to the deviation of that individual's RPB-2 abundance from the day 0 population average.

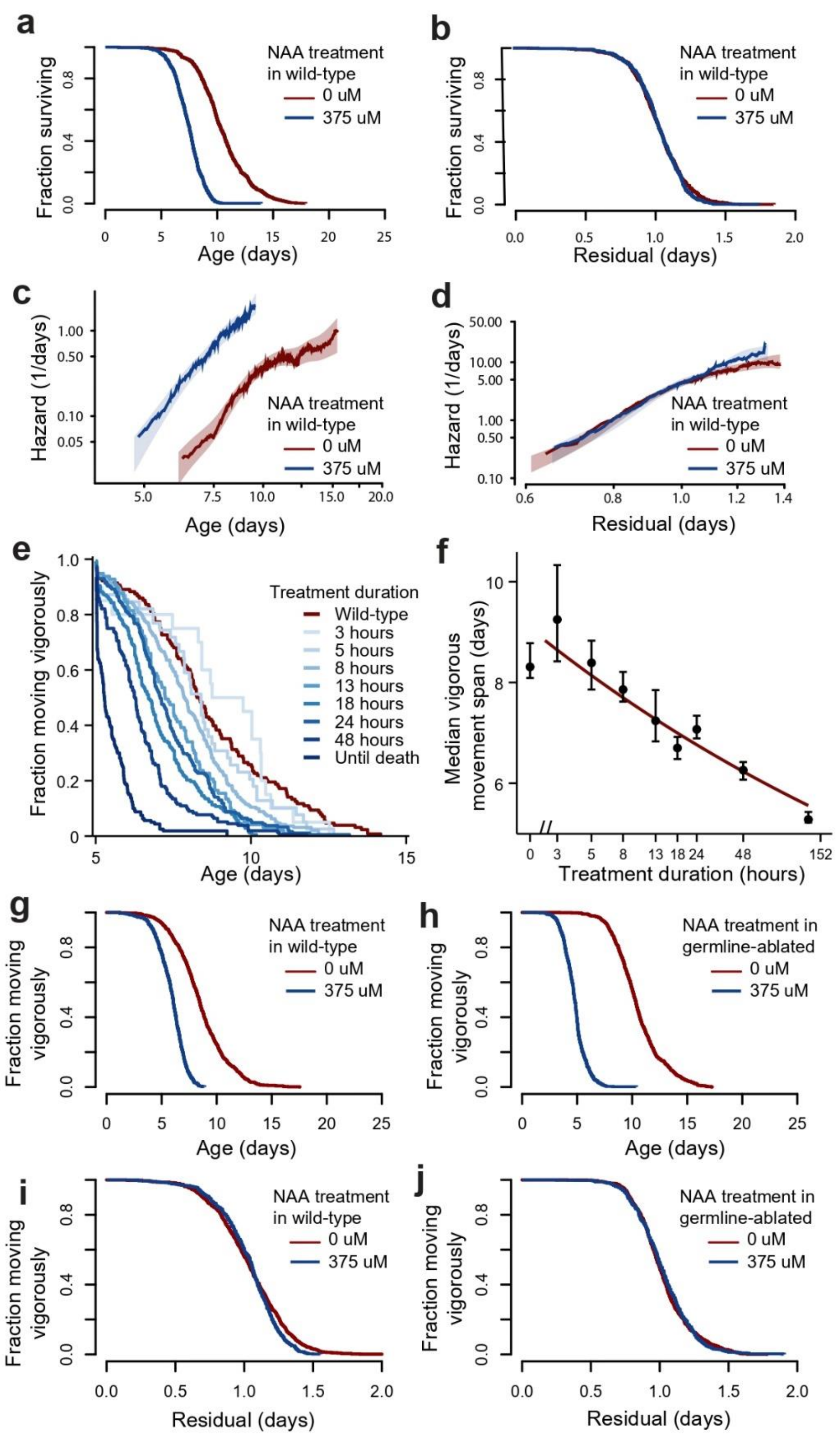


**Figure S4—Transient increases in RNAPolII degradation rates permanently limit total mRNA abundance and lifespan via accelerated aging. a.** For wild-type populations (N2), a Kaplan-Meier analysis comparing the effect on lifespan of NAA-induced life-long, high RPB-2 degradation starting in young adulthood (day 2) (*blue,* n=600) compared to unperturbed populations (*red,* n=919). **b.** The residuals of an accelerated-failure time (AFT) model fit to the data in *a*, highlighting any deviations from temporal scaling (no significance, *P*>0.2). **c.** The same data in *a*, plotted as hazard curve estimates of the trajectory in the risk of death. **d.** The same AFT residuals as in *b,* but replotted as hazard curves. **e.** The effect of transient increases in RPB-2 degradation, as in Fig. 4c, but considering individuals' remaining "healthspan", defined as the time until age-associated vigorous movement cessation (VMC) (n=1668). **f.** Quantification of *e*, comparing the relationship between median VMC and the duration of high RPB-2 degradation treatment, fit with a linear regression line. **g.** The effect on VMC of NAA-induced life-long, high RPB-2 degradation in wild-type from the populations in *a*, and in **h.**, germline-ablated *glp-1(e2141)* populations (*blue,* n=636; *red,* n=1061). **i-j.** The residuals of AFT models fit for VMC accounting for differences in timescale of the populations shown in *g* (KS *P*=0.00024) and *h* (no significance, KS *P*=0.029), respectively.

### Supplemental Note 1

Estimation of absolute mRNA abundance using RNA-Seq data with ERCC spike-ins

#### 1.1 Cluster-competition model and the naïve transcriptome-size formula

We seek an estimator of the per-sample absolute mRNA abundance using ERCC spike-ins as an external molar reference. For sample $s$, let $L_s$ denote the total cDNA molarity entering the sequencing lane, $E_s$ the ERCC component and $M_s$ the endogenous mRNA component, so that

$$L_s = E_s + M_s. \tag{1}$$

Because mRNA and ERCC cDNA molecules are sequenced together on the same flow cell, they compete for a finite number of clusters. The observed counts $\hat{L}_s = \hat{E}_s + \hat{M}_s$ therefore reflect not the absolute molarities but their fractional shares of the library, weighted by the total cluster allocation $\alpha_s$ assigned to sample $s$:

$$\hat{M}_s = \alpha_s \cdot \frac{M_s}{E_s + M_s}, \qquad \hat{E}_s = \alpha_s \cdot \frac{E_s}{E_s + M_s}. \tag{2}$$

For an individual ERCC species $i$ with input molarity $E_{is}$ (so $\sum_i E_{is} = E_s$ and $\sum_i \hat{E}_{is} = \hat{E}_s$),

$$\hat{E}_{is} = \frac{\alpha_s}{E_s + M_s} E_{is}. \tag{3}$$

Dividing (2) by (3) cancels both $\alpha_s$ and $E_s + M_s$,

$$\frac{\hat{M}_s}{\hat{E}_{is}} = \frac{M_s}{E_{is}}, \tag{4}$$

which is the central identity: per sample, the ratio of observed mRNA to observed ERCC is equal to the ratio of their absolute molarities. Defining

$$\beta_s \equiv \frac{\hat{M}_s}{M_s} \tag{5}$$

as the per-sample scaling between observed and absolute mRNA, (4) becomes $\hat{E}_{is} = \beta_s \cdot E_{is}$, i.e. observed ERCC counts are linear in their known input concentrations through the same $\beta_s$ that links observed and absolute mRNA. The absolute mRNA abundance of sample $s$ — its transcriptome size — is therefore

$$\mathrm{TS}_s \equiv M_s = \frac{1}{\beta_s} \sum_j \hat{M}_{js}. \tag{6}$$

Equation (6) is well defined but is unsatisfactory in two practical respects. First, the numerator $\sum_j \hat{M}_{js}$ is dominated by a small number of highly abundant transcripts —for example, in gravid *C. elegans*, vitellogenin mRNAs alone can constitute $> 40\%$ of the polyadenylated transcriptome. If we assume that sequencing errors produce fold-change effects, then the errors in highly expressed genes will contribute disproportionately. Second, $\beta_s$ estimated from a raw-scale fit of (3) is computed based on the abundance of ERCC transcripts, whose concentrations within each ERCC library span ~6 orders of magnitude and therefore again will be dominated by the small number of very high abundance ERCC transcripts. Both limitations motivate a regression-based reformulation in which the numerator and denominator of (6) are estimated jointly on the log scale from the full mRNA and ERCC count distributions.

##### 1.2 Dual-regression decomposition

We re-express the transcriptome-size estimator as a ratio of two regression-derived quantities,

$$\mathrm{TS}_s = \frac{\lambda_s}{d_s}, \tag{7}$$

where $\lambda_s$ is an mRNA-derived sample-specific scale factor that aligns the *full* mRNA count distribution of sample $s$ to that of a designated reference sample $r$, and $d_s$ is an ERCC-derived sample-specific depth factor that captures the technical scaling shared across all ERCC species in sample $s$. Both factors are estimated on the log scale by Gaussian linear models and back-transformed by the delta method; this yields per-sample standard errors and per-feature residual diagnostics for each component of the ratio.

Throughout, $g \in \{1, \dots, G\}$ indexes retained mRNA genes, $k \in \{1, \dots, K\}$ indexes retained ERCC species, $s \in \{1, \dots, S\}$ indexes samples passing filtering (see Methods), and $Y_{gs}, R_{ks} \in \mathbb{N}_0$ denote the observed mRNA and ERCC counts. The known ERCC input concentrations,$c_k$ are all greater than zero (attomoles $\cdot$ µL$^{-1}$). We designate one sample $r$ — the first sample of the youngest (or untreated) reference group — as the reference, and fix $\lambda_r = 1$.

##### 1.3 Numerator: pooled regression for the mRNA scale factor $\lambda_s$

The working biological hypothesis is that, to first order, mRNA distributions between samples differ predominantly by a global multiplicative scale rather than by complete redistribution of gene-specific abundances. On the log scale, this corresponds to an additive sample shift:

$$\log Y_{gs} = \log Y_{gr} + \alpha_s + \epsilon_{gs}, \qquad \alpha_r = 0, \tag{8}$$

with $\epsilon_{gs}$ a zero-centred residual capturing per-gene departure from pure scaling. The multiplicative mRNA scale factor is then

$$\lambda_s \equiv \exp(\alpha_s). \tag{9}$$

Rather than fitting (8) one sample at a time against the reference, we pool all genes and samples into a single long table $\{(g, s, Y_{gs})\}$ and fit the Gaussian linear model

$$\log(Y_{gs}) = \beta_0 + \alpha_s + \epsilon_{gs}, \qquad \alpha_r = 0, \tag{10}$$

with the reference sample absorbed into the intercept ($\beta_0$ represents the reference-sample log location). The fit is performed using `RcppArmadillo::fastLm`. The pseudocount of 1 is retained for numerical robustness in residual computations but has negligible effect on the coefficient estimates because genes retained after filtering satisfy $Y_{gs} \geq 10$ in $\geq 95\%$ of samples, so that $\log(Y_{gs} + 1) \approx \log Y_{gs}$.

The estimator of the mRNA scale factor is

$$\hat{\lambda}_s = \exp(\hat{\alpha}_s), \tag{11}$$

with delta-method standard error

$$\mathrm{SE}(\hat{\lambda}_s) \approx \left|\tfrac{\mathrm{d}}{\mathrm{d}\alpha} \exp(\alpha)\right|_{\alpha=\hat{\alpha}_s} \cdot \mathrm{SE}(\hat{\alpha}_s) = \hat{\lambda}_s \cdot \mathrm{SE}(\hat{\alpha}_s). \tag{12}$$

Because $\hat{\alpha}_s$ is jointly informed by all $G$ retained genes, a single dominant transcript contributes only one of $G$ observations to its estimation, so $\hat{\lambda}_s$ is substantially more robust to highly expressed outlier genes than the raw mRNA sum $\sum_j \hat{M}_{js}$ that appears in (6). Per-gene residuals

$$\hat{\epsilon}_{gs} = \log Y_{gs} - \log \hat{Y}_{gs} \tag{13}$$

quantify deviation from the pure-scaling approximation. We assess (8) empirically by comparing the residual empirical cumulative distribution functions across age groups (Fig. 1e); the unimodal and approximately symmetric $\log_2$ fold-change distributions of Fig. 1b indicate that, in the regimes studied here, age-related mRNA changes are dominated by a multiplicative global shift rather than by wholesale redistribution across genes, and KS-type effect sizes between young and old residual ECDFs are small (Fig. 1e).

##### 1.4 Denominator: per-sample offset regression for the ERCC depth factor $d_s$

Within a single sample $s$, we assume that every ERCC species experiences the same library preparation efficiency, capture efficiency, sequencing depth, and cluster-competition environment — quantities we collectively encode in a per-sample technical depth factor $d_s$. The expected ERCC counts therefore satisfy

$$\mathbb{E}[R_{ks}] = d_s \cdot c_k, \tag{14}$$

so that on the log scale

$$\log \mathbb{E}[R_{ks}] = \log d_s + \log c_k, \tag{15}$$

which is a one-parameter regression problem per sample with $\log d_s$ as the only unknown coefficient. We fit it as a sample-by-sample Gaussian linear model with an offset:

$$\log(R_{ks}) = \delta_s + \log(c_k) + \epsilon_{ks}, \tag{16}$$

equivalently $\log(R_{ks}) \sim 1 + \text{offset}\big(\log(c_k)\big)$. The estimator of the depth factor is

$$\hat{d}_s = \exp(\hat{\delta}_s), \tag{17}$$

with delta-method standard error

$$\text{SE}(\hat{d}_s) \approx \hat{d}_s \cdot \text{SE}(\hat{\delta}_s). \tag{18}$$

Per-species residuals

$$\hat{\epsilon}_{ks} = \log R_{ks} - \big(\hat{\delta}_s + \log c_k\big) \tag{19}$$

provide diagnostics for departure from the linear log–log relationship. Samples or ERCC species displaying systematic non-linearity — saturation at the high-concentration end or drop-out at the low-concentration end — are identified directly from (19); we excluded ERCC species with $c_k$ outside the linear range ($c_k \in [0.5,\ 10^4]$ (attomoles $\cdot$ µL$^{-1}$) prior to fitting the mouse pseudobulks (Fig. 1g) where dynamic-range compression was apparent. Formulation (16) uses all $K$ ERCC species jointly, preserving per-species information that would be lost if $d_s$ were estimated from $\sum_k R_{ks} / \sum_k c_k$.

##### 1.5 Combining numerator and denominator: conditional independence

The transcriptome-size estimator is

$$\widehat{\text{TS}}_s = \frac{\hat{\lambda}_s}{\hat{d}_s} = \exp(\hat{\alpha}_s - \hat{\delta}_s), \tag{20}$$

and on the log scale takes the particularly clean form

$$\log \widehat{\mathrm{TS}}_s = \hat{\alpha}_s - \hat{\delta}_s, \tag{21}$$

i.e. a difference of two regression coefficients obtained from disjoint subsets of the data: $\hat{\alpha}_s$ is informed only by the mRNA count matrix $\{Y_{gs}\}$ and $\hat{\delta}_s$ only by the ERCC count matrix $\{R_{ks}\}$. Here we take the assumption that, conditional on these two count matrices, $\hat{\alpha}_s$ and $\hat{\delta}_s$ are mutually uninformative — knowing the residuals of one regression contributes nothing to the residuals of the other — and we therefore treat them as approximately independent for the purpose of uncertainty propagation,

$$\mathrm{Cov}(\hat{\lambda}_s, \hat{d}_s) \approx 0. \tag{22}$$

This is a working approximation rather than a strict identity: at the data-generating level, both regressions are conditional on the unobserved cluster-allocation factor $\alpha_s$ of equation (2), which in principle ties them together upstream of the observed counts. The independence in (22) refers to the conditional independence of the two estimators given the count data actually used to fit them.

#### 1.6 Uncertainty propagation by the delta method

Under (22), the raw-scale delta-method variance of $\widehat{\mathrm{TS}}_s = f(\hat{\lambda}_s, \hat{d}_s) = \hat{\lambda}_s/\hat{d}_s$ follows from

$$\frac{\partial f}{\partial \lambda} = \frac{1}{d}, \qquad \frac{\partial f}{\partial d} = -\frac{\lambda}{d^2}, \tag{23}$$

giving

$$\mathrm{Var}(\widehat{\mathrm{TS}}_s) \approx \frac{\mathrm{Var}(\hat{\lambda}_s)}{\hat{d}_s^2} + \frac{\hat{\lambda}_s^2}{\hat{d}_s^4}\,\mathrm{Var}(\hat{d}_s), \tag{24}$$

and

$$\mathrm{SE}(\widehat{\mathrm{TS}}_s) \approx \sqrt{\left(\frac{\mathrm{SE}(\hat{\lambda}_s)}{\hat{d}_s}\right)^2 + \left(\frac{\hat{\lambda}_s \cdot \mathrm{SE}(\hat{d}_s)}{\hat{d}_s^2}\right)^2}. \tag{25}$$

The symmetric raw-scale 95% confidence interval reported in the figures is

$$\widehat{\mathrm{TS}}_s \pm 1.96\ \mathrm{SE}(\widehat{\mathrm{TS}}_s). \tag{26}$$

An equivalent log-scale formulation exploits the additive structure of (21). Under (22),

$$\mathrm{Var}(\log \widehat{\mathrm{TS}}_s) \approx \mathrm{Var}(\hat{\alpha}_s) + \mathrm{Var}(\hat{\delta}_s), \tag{27}$$

and back-transformation yields a strictly positive multiplicative interval,

$$\Big[\widehat{\mathrm{TS}}_s \cdot \exp\!\big(-1.96\ \mathrm{SE}(\log \widehat{\mathrm{TS}}_s)\big),\ \widehat{\mathrm{TS}}_s \cdot \exp\!\big(+1.96\ \mathrm{SE}(\log \widehat{\mathrm{TS}}_s)\big)\Big]. \tag{28}$$

The raw-scale (26) and log-scale (28) intervals are indistinguishable at the magnitudes of $\widehat{\mathrm{TS}}_s$ encountered in our datasets (relative TS $\gtrsim 0.05$); we report (26) for direct comparability with the raw-scale axes used in the figures. The log-scale form is preferable in any setting where $\widehat{\mathrm{TS}}_s$ approaches zero, because (26) can otherwise produce negative lower bounds. Both (26) and (28) are first-order Wald approximations and would become anti-conservative if $\hat{\alpha}_s$ or $\hat{\delta}_s$ had strongly skewed or heavy-tailed sampling distributions — for example in samples retaining only a small number of ERCC species after filtering. Where finite-sample exactness is required, bootstrapping over genes (for $\hat{\lambda}_s$) or ERCC species (for $\hat{d}_s$) provides a straightforward extension.

#### 1.7 Relationship to existing normalisation schemes

Standard RNA-seq normalisation methods are designed to remove per-sample technical scaling so that genes can be compared across samples on a common relative scale; by construction, they discard the very quantity that absolute transcriptome size aims to estimate.

*Library-size normalisations* such as CPM, RPKM and TPM divide counts by the total mRNA count of the sample, forcing each sample's estimated transcriptome size to be constant by construction. Global mRNA shifts are therefore invisible to any analysis built on these counts.

*Median-of-ratios estimators* (DESeq2) and *deconvolution-based size factors* (`scran::calculateSumFactors`) conventionally identify a per-sample scaling factor under the working assumption that most genes are not differentially expressed between samples. The factor they estimate is, up to a constant, related to our $\lambda_s$, but its intended use is to divide it out of the counts — again removing the global signal. Furthermore, these methods lack a denominator anchored to a physical molar reference, and so cannot in principle recover absolute transcriptome size: they identify the multiplicative shift between samples but cannot say which sample, if any, is at "absolute" scale.

*Spike-in summation*, in which $d_s$ is estimated from $\sum_k R_{ks}$ alone (or equivalently from the ratio $\sum_k R_{ks} / \sum_k c_k$, as in `scran::computeSpikeFactors`), discards per-species information and is sensitive to saturating or drop-out ERCCs. The regression-based $\hat{d}_s$ of (17) has comparable mean behaviour but tighter standard errors at fixed $K$ and exposes outlier ERCC species through the residuals (19).

The estimator $\widehat{\mathrm{TS}}_s = \hat{\lambda}_s / \hat{d}_s$ therefore occupies a different niche from these methods: the numerator preserves the global mRNA shift that standard normalisations remove, and the denominator anchors that shift to a known molar reference through the ERCC spike-ins, yielding an estimate in absolute (rather than relative) transcriptome units with calibrated per-sample uncertainty.

### Supplementary Note 2

### Estimating protein synthesis and degradation rate constants *in vivo* using quantitative imaging data

#### 1. Estimating the RPB-2 degradation constant

**Estimating the size of the un-degraded fraction:** From microscopy images, we obtained the total nuclear intensity of each individual in three conditions: “Unconverted” individuals not exposed to photoconversion, the same individuals immediately (< 1 hour) after photoconversion of the dendra2 fluorophore, and the same individuals a third time after 24 hours of recovery after photoconversion. “Unconverted” and “converted” individuals were collected on days 0, 1, and 4 of adulthood, and “recovered” individuals were collected on days 1, 2, and 5.

“Unconverted” individuals were not used in the calculation of degradation rate. Instead, they were used as a negative control to validate assumptions about image analysis background correction strategies, and to quantify a negligible contribution of autofluorescence.

To measure the degradation rate, we calculate the ratio

$$\log_2 f = \log_2 \frac{1}{N_R} \sum_j I_{R,j} - \log_2 \frac{1}{N_P} \sum_j I_{P,j}$$

which quantifies the average intensity of “recovered” individuals $I_R$ relative to the average intensity of “just photoconverted” $I_P$. An *f* value of 1 indicates no degradation of the photoconverted RPB-2 occurred during the recovery period, and an f value of 0 indicates all photoconverted RPB-2 was degraded. We made sure that all *f* values took an intermediate value, as very small *f*s would indicate that we chose a recovery period too long to accurately estimate the degradation rate constant.

Population averages were used because, though the population $I_R$include the same individuals as in $I_P$, we were unable to match the identity of each individual $I_{R,j}$ to its previous measurement in $I_{P,j}$ because individual’s positions were randomized and identities lost during the recovery period.

**Batch Correction:** To obtain large population sizes $N_R$ and $N_P$, we must aggregate image data collected in multiple separate biological replicates, with population sizes $N_R^1, N_R^2, N_R^3, \ldots, N_R^K$ . For our final estimate of $k_d$ we use the “batch-corrected” ratio $f$ defined as the weighted average of all replicates; $f^k$ values:

$$f^{all} = \frac{\sum_k \left(N_R^k + N_P^K\right) f^k}{\sum_k \left(N_R^k + N_P^K\right)}$$

**Estimating the degradation rate constant:** The degradation rate constant is defined as parameter governing the first-order degradation of RPB-2 $dR/_{dt} = -k_d R$ . Note that we are measuring only the photoconverted fraction, for which no de novo synthesis occurs. The solution for these

dynamics is $R(t) = c_1\, e^{-k_d t}$, an exponential decay from the initial condition $R(0) = c_1$ to $R(\infty) = 0$
.

Using measurements of R at two timepoints, first at the "just photoconverted" timepoint $t = 0$ and again in the "recovered" timepoint $t = t_1$, we can solve for the degradation rate constant

$$k_d = \frac{1}{t_1} \ln \frac{R(t=0)}{R(t=t_1)}$$

which we can write in terms of degradation fraction

$$k_d = -\frac{1}{t_1} \ln f$$

To obtain confidence intervals for $k_d$, we consider $k_d$values estimated across bootstrap samples (with replacement) selected from the set of all individuals $\{I_{R,j}, I_{P,j}\}$.

**Quantifying changes in the degradation rate constant over time:** To estimate changes in the bootstrap samples during aging, we look at the ratio of $k_d$ at two different ages

$$k_d^{old} \Big/ k_d^{young} = -\frac{1}{t_2} \ln f^{old} \Big/ -\frac{1}{t_1} \ln f^{young}$$

We can then use the same bootstrapping procedure for estimating confidence intervals on this ratio.

#### 2. Estimating the RPB-2 synthesis rate

**Estimating the size of the newly synthesized fraction:** We follow the same experimental design as before, but this time comparing untreated individuals, the same individuals immediately after photobleaching $I_{PB}$, and the same animals a third time after 24 hours of recovery, $I_{RE}$the and the same individuals following

Again, we use the "unconverted" individuals only to confirm full photoconversion and confirm that autofluorescence does not influence our other measurements. The mean intensity of the newly synthesized fraction of RPB-2 during the recovery period is simply

$$S = \frac{1}{N_R} \sum_j I_{RE,j}$$

**Estimating the synthesis rate:** During the recovery period, individuals are simultaneously synthesizing new protein and degrading both the "old" photo-bleached and newly-synthesized proteins. We cannot inhibit protein degradation for the full 24 hour recovery period without causing many unwanted side effects that would likely cause protein synthesis rates to deviate from their physiologic levels. So, instead, we correct our newly synthesis fraction for the effects of protein degradation, whose effects we can predict using our knowledge of the degradation rate constant $k_d$.

Adding synthesis to the same kinetics as before

$$dR/_{dt} = k_s - k_d R$$

Because we are able to photobleach the entire population of R so that $R(t = 0) = 0$ , we can obtain the standard solution

$$R(t) = \frac{k_s}{k_d}\left(1 - e^{-k_d t}\right)$$

We can then use our measurement of the newly synthesized fraction $R(t = t_1) = S$ to solve for $k_s$

$$k_s = S\left(\frac{k_d}{(1 - e^{-k_d t_1})}\right)$$

Our procedure is therefore equivalent to as our measurement of RPB-2 intensity by a “correction factor” that accounts for the cumulative effect of degradation. For example, during youth we find $k_d$ takes a value of approximately 0.75, and so after one day of recovery, $t_1 = 1$, $k_s$is 1.4 times larger than S, indicating that our raw measurement is 1/1.4 = 70% of the level we’d have measured in the absence of any RPB-2 degradation.

We can then estimate confidence intervals for the synthesis rate, and for changes in the synthesis rate, by estimating $k_s$ on bootstrap samples taken from the population of all individuals measured in both photobleaching and photoconversion experiments $\{I_{RE,j}, I_{R,j}, I_{P,j}\}$.

**Estimating the equilibrium set point**

The equilibrium set point is the value of R at which $dR/_{dt} = 0$ and the system is at steady state. We can estimate this value as $k_s/_{k_d}$, again with confidence intervals obtained from estimating the ratio across bootstrap samples drawn from the population of all individuals $\{I_{RE,j}, I_{R,j}, I_{P,j}\}$.

**Estimating changes in the equilibrium set point**

We can finally estimate the magnitude of age-associated changes in the equilibrium set point as

$$\Delta Eq = \frac{k_s^{old}/_{k_d^{old}}}{k_s^{young}/_{k_d^{young}}}$$

We can obtain confidence intervals on changes in the equilibrium set point via bootstrapping over our entire corpus of image data, $\{I_{RE,j}, I_{R,j}, I_{P,j}\}$ measured in young individuals and also $\{I_{RE,j}, I_{R,j}, I_{P,j}\}$ measured in old individuals.
